# Hypersensitivity to uncertainty is key feature of subjective cognitive impairment

**DOI:** 10.1101/2021.12.23.473986

**Authors:** Bahaaeddin Attaallah, Pierre Petitet, Elista Slavkova, Vicky Turner, Youssuf Saleh, Sanjay G. Manohar, Masud Husain

## Abstract

With an increasingly ageing global population, more people are presenting with concerns about their cognitive function, but not all have an underlying neurodegenerative diagnosis. Subjective cognitive impairment (SCI) is a common condition describing self-reported deficits in cognition without objective evidence of cognitive impairment. Many individuals with SCI suffer from depression and anxiety, which have been hypothesised to account for their cognitive complaints. Despite this association between SCI and affective features, the cognitive and brain mechanisms underlying SCI are poorly understood. Here, we show that people with SCI are hypersensitive to uncertainty and that this might be a key mechanism accounting for their affective burden. Twenty-seven individuals with SCI performed an information sampling task, where they could actively gather information prior to decisions. Across different conditions, SCI participants sampled faster and obtained more information than matched controls to resolve uncertainty. Remarkably, despite their ‘urgent’ sampling behaviour, SCI participants were able to maintain their efficiency. Hypersensitivity to uncertainty indexed by this sampling behaviour correlated with the severity of affective burden including depression and anxiety. Analysis of MRI resting functional connectivity revealed that both uncertainty hypersensitivity and affective burden were associated with stronger insular-hippocampal connectivity. These results suggest that altered uncertainty processing is a key mechanism underlying the psycho-cognitive manifestations in SCI and implicate a specific brain network target for future treatment.

## Introduction

With an ageing population, an increasing number of people are seeking medical advice for concerns about cognitive decline (Deary et al., 2009; Harada et al., 2013). While in some individuals these complaints might be related to a progressive pathological process such as Alzheimer’s disease, they can also be expressed by people without an underlying neurodegenerative disorder (McWhirter et al., 2020). When objective clinical evidence of significant cognitive impairment is not evident alongside self-reported cognitive complaints, individuals are categorised as having subjective cognitive impairment/decline (SCI/D) (Jessen et al., 2014, 2020; Reid and MacLullich, 2006). Although most follow a relatively benign course, a small proportion develops objective cognitive impairment and subsequently progress to dementia (Arvanitakis et al., 2018; Kryscio et al., 2014; Mendonça et al., 2016). Nevertheless, it remains unclear what drives subjective cognitive complaints in those people who do not have evidence of underlying neurodegeneration. Understanding the mechanisms of cognitive, behavioural, and psychiatric manifestations in SCI is thus crucial to improve clinical outcomes and enhance understanding of presentation of people who present with cognitive concerns.

A wealth of evidence suggests a particularly high prevalence of a range of mental health problems associated with SCI, in particular affective disorders such as anxiety and depression (Hill et al., 2016; Hohman et al., 2011; Pavisic et al., 2021; Reid and MacLullich, 2006). Similarly, people who primarily suffer from these psychiatric disorders often report sub-optimal cognitive performance, further emphasising the intertwined relationship between affective burden and subjective cognitive experience (Millan et al., 2012). Moreover, treating anxiety and depression may improve subjective cognitive complaints in individuals with SCI (Allott et al., 2020). Despite the association between SCI and affective burden being increasingly recognised, little is understood about the underlying cognitive mechanisms and brain networks involved.

A rich body of theoretical and empirical work suggests that affective dysregulation might be related to uncertainty processing and related behaviours (Bishop and Gagne, 2018; Carleton, 2016; Grupe and Nitschke, 2013; Gu et al., 2020). People who express higher levels of anxiety and depression often report higher levels of intolerance to uncertainty (Boelen et al., 2016; Boswell et al., 2013; Carleton et al., 2012; McEvoy and Mahoney, 2011; Saulnier et al., 2019). Mechanistically, intolerance to uncertainty might be reflected in several cognitive and behavioural processes underpinning goal-directed behaviour when people decide and act under uncertainty (Grupe and Nitschke, 2013). For example, when someone is crossing the road, they make their decision based on how confident they are that the environment is safe (i.e., they have an assessment of how uncertain the environment is for their intended action) (Bach and Dolan, 2012; Gottlieb and Oudeyer, 2018). If uncertainty is high, agents often try to reduce it by gathering information to inform their decision (e.g., checking passing cars and traffic lights and looking for a safer place to cross). People who are more sensitive to uncertainty might have an exaggerated estimation of uncertainty or preparedness when required to face it, eventually affecting their decisions and outcomes (Grupe and Nitschke, 2013). Similarly, uncertainty sensitivity might affect self evaluation of cognitive abilities (e.g., having lower confidence in recollection) amplifying memory complaints and subsequent emotional reaction (Fitzgerald et al., 2017; Nelson, 1990). Such a framework, which involves estimation, valuation, preparation, and learning under uncertainty allows a detailed investigation of the psychopathology of affective dysfunction (Gottlieb and Oudeyer, 2018; Grupe and Nitschke, 2013; Sharot and Sunstein, 2020).

Investigation of the dynamics of how people decide and act under uncertainty might hold an important key to understanding the relationship between SCI and affective dysfunction. This might be challenging to achieve using classical behavioural paradigms, e.g., beads task or variants of it (Phillips et al., 1966). These paradigms often involve randomly drawing samples from a distribution to make inferences about the distribution (e.g., deciding the predominant colour of beads in a jar). However, when people gather information to reduce uncertainty, they dynamically assess their environment and update their expectations in order to decide whether a new piece of evidence is needed and whether they can tolerate its cost (Juni et al., 2016; Petitet et al., 2021). While the economic aspect of this behaviour has been extensively examined in previous studies (Clark et al., 2006; Jones et al., 2019; Juni et al., 2016), investigation into *how* information is gathered is limited. Capturing behavioural markers that might not be directly apparent using such tasks (e.g., sampling speed and efficiency) might provide important insights into underpinning mechanisms of affective disorders. This distinction has been formalised as ‘active’ information gathering, characterised by situations in which participants have agency over not only how much information they gather but also how information is collected in face of uncertainty (e.g., what resources to consult and when) (Gottlieb and Oudeyer, 2018; Petitet et al., 2021).

In this study, we adopted this approach using a recently developed behavioural paradigm to investigate how people with SCI decide and act (gather information) under uncertainty (Petitet et al., 2021). A crucial question of this study was whether uncertainty processing is associated with affective burden. Further, to investigate the underlying brain structures and networks that might be implicated in the process, brain resting functional neuroimaging (rfMRI) data were also collected. In an *active form* of the task, participants collected informative clues, which came at a known cost, to reduce their uncertainty before committing to decisions. Crucially, they were allowed to freely gather information whenever and in whichever way they wanted. In a *passive form* of the task, this agency over uncertainty was removed. Participants were allowed only to accept or reject offers that had fixed levels of uncertainty weighed against potential reward. Decisions were made based on whether tolerating uncertainty was worth the reward on offer. This enabled us to calculate how people weigh uncertainty against reward in a passive environment where agency over uncertainty is absent. Before decisions, participants also reported their subjective uncertainty, enabling us to measure the accuracy of uncertainty estimation that might influence both active and passive behaviour.

The results from the behavioural tasks showed that individuals with SCI gather significantly more information than healthy matched controls before they commit to final decisions. They did this regardless of the cost of information and at a faster rate than controls. Despite this faster sampling, SCI participants impressively were capable of maintaining their sampling efficiency (i.e., gathering samples that were as informative as controls). This meant that they exceeded the speed-efficiency trade-off that characterises sampling behaviour of healthy controls. Crucially, in individuals with SCI, sampling speed and over-sampling (indices of heightened sensitivity to uncertainty) were associated with affective burden (derived from self-report measures of anxiety and depression).

By contrast, when they had no agency over the reward and uncertainty on offer (passive choice task), SCI participants had intact metacognitive assessment and valuation of uncertainty. This suggests that controllability when dealing with uncertainty might be a crucial aspect in affective dysfunction, as differences between SCI participants and controls were apparent only when they had agency over uncertainty (i.e., during the active information gathering phase preceding decisions).

Functional neuroimaging analysis investigating whole-brain resting connectivity between regions of interest across all known brain networks revealed that individuals with SCI, in comparison to healthy controls, had increased insular hippocampal connectivity. Further, the strength of this connectivity in SCI correlated with sensitivity to uncertainty indexed by sampling speed. Across all study participants, insular-hippocampal connectivity was also found to correlate with reported affective burden. This suggested that urgent sampling behaviour when faced with uncertainty might mediate the association between heightened insular-hippocampal connectivity and affective dysfunction, a findings which was confirmed with Bayesian mediation analysis.

Taken together, the results indicate that hyper-reactivity to uncertainty might be a key mechanism in SCI, and link this process to insular cortex and hippocampus.

## Results

### Experimental design

Participants performed a recently developed behavioural task (Petitet et al., 2021) designed to investigate active information gathering and decision making under uncertainty (Figure 1). In this paradigm, participants were asked to maximise their reward by trying to localise a hidden purple circle of a fixed size as precisely as possible. They could reduce uncertainty about the location of the hidden circle by touching the screen at different locations to obtain informative clues: if a purple dot appeared where they touched, this meant that the location was situated inside the hidden circle, otherwise, the dots were coloured white. Obtaining these clues came at a cost (*η*_*s*_) that participants had to pay from an initial credit reserve (*R*_0_) they started each trial with. Participants could sample the search field freely without constraints to the location or the speed at which they touch the screen. At the end of each trial, participants were required to move a blue disc to where they thought the hidden circle was located. A feedback was given after this, indicating how many credits participants won (or lost) based on how precise their localisation was and the credits they lost to obtain information (i.e., participants had to make a trade-off between obtaining more information and the cost of this information). There were two levels of sampling cost (low and high) and two levels of initial credit reserve (low and high). Uncertainty in the task was quantified as expected error (*EE*) which is the average error that an optimal agent is expected to obtain when placing the blue disc at the best possible location. For more details see *Methods* and Supplementary information.

**Figure 1:**
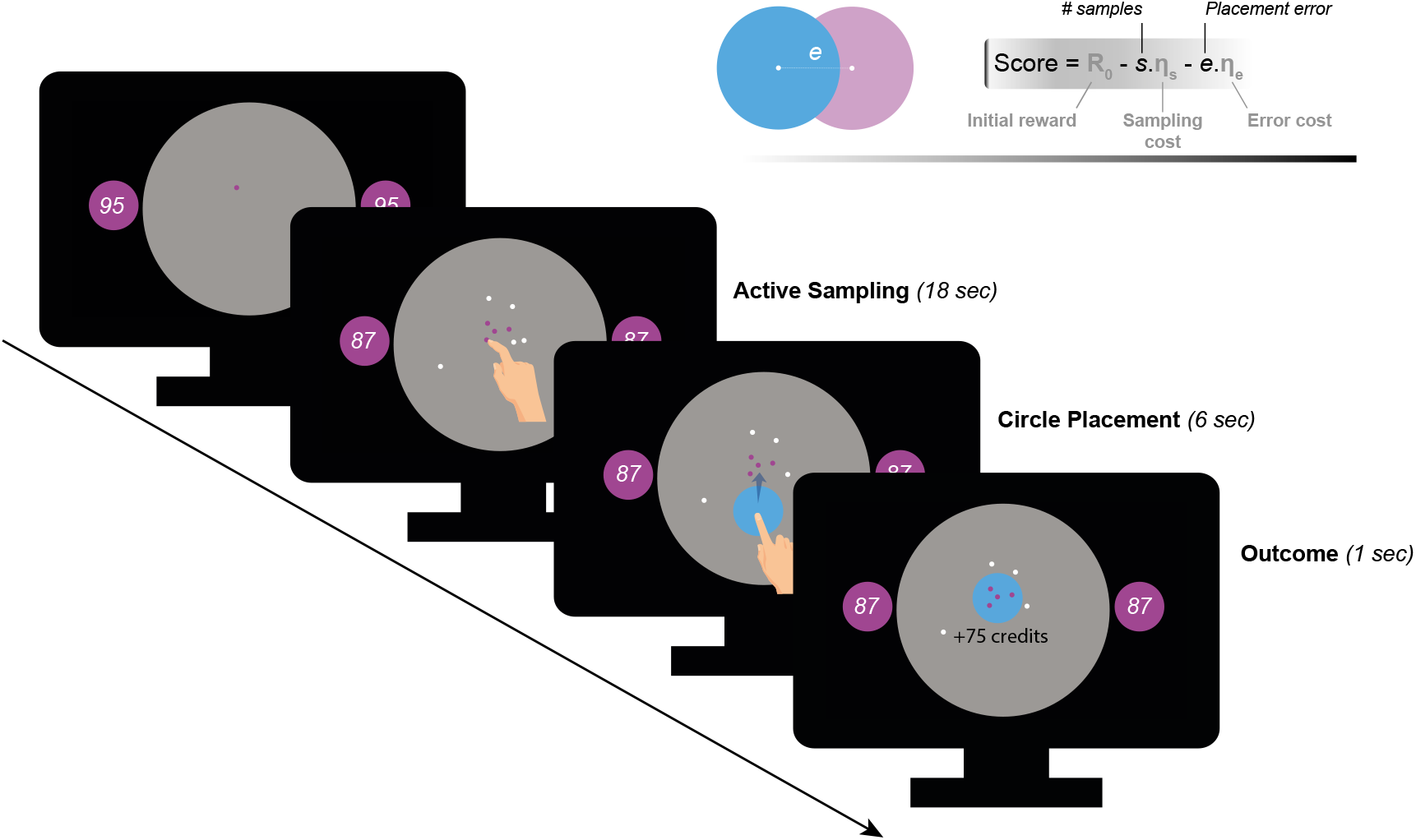
Task paradigm – Active information sampling. Participants were required to find the location of a hidden purple circle as precisely as possible. Clues about the location of the hidden circles could be obtained by touching the screen at different locations. This yielded dots that were coloured either purple or white depending on whether they were situated inside or outside the hidden circle: purple dots were inside and white dots were outside. Two circles of the same size as the hidden circle were always on display on either side of the screen to limit memory requirements of the task. Inside these two circles, an initial credit reserve (*R*_0_) was displayed. There were two levels of *R*_0_: low = 95 credits and high = 130 credits. At the beginning of each trial, a purple dot was always shown to limit initial random sampling. Participants then had 18 seconds on each trial, during which they could sample without restrictions to speed, location or number of samples. Each time they touched the screen to add a dot, the number of credits available decreased depending on the cost of sampling (*η*_*s*_) on that trial. There were two levels of *η*_*s*_: low = −1 credit/sample and high = −5 credits/sample. Once the 18 seconds have passed, a blue disc of the same size as the hidden circle appeared at the centre of the search field. Participants were then required to drag this disc on top of where they thought the hidden circle was located. Following this, the score they obtained on that trial was calculated and presented as feedback at the end of the trial.

### Demographics

All participants (healthy controls and individuals with SCI) had ACE-III cognitive scores within normal performance limits (> 87/100) (Bruno and Vignaga, 2019; Elamin et al., 2016; Hsieh et al., 2013). There was no significant difference between SCI participants and controls in cognitive scores (Controls: *μ* = 97.89, *SD* = 1.80; SCI: *μ* = 95.41, *SD* = 4.21; *z* = 1.91, *p >* 0.05). Consistent with previous reports (Hill et al., 2016; Hohman et al., 2011; Pavisic et al., 2021; Reid and MacLullich, 2006), SCI participants in the study were significantly more depressed and anxious than healthy controls (Depression: *z* = 4.41, *p* < 0.001, Anxiety: *z* = 3.08, *p* < 0.01; Table 1 & Figure 5a.). Since depression and anxiety correlated positively with each other (Spearman’s *R*^2^ = 0.45, *p* < 0.001, Figure 5b.), a principal component analysis (PCA) was performed to extract a dimension that accounts for the maximum shared variance between the two measures. This dimension could be regarded as a measure of affective burden in participants and accounted for 84% of the variance shared between depression and anxiety. Higher scores of affective burden indicate more severe depression and anxiety. There was no significant correlation between cognitive scores and this affective dimension (*p >* 0.05, controlling for age and gender).

**Table 1:** Demographics. ACE-III: Addenbrooke’s Cognitive Examination. BDI-II: Beck Depression Inventory. HADS: Hospital Anxiety Depression Scale. * Student-test or Wilcoxon rank-sum test for parametric and non-parametric data, respectively.

| N (M/F) | Controls |  | SCI |  | p-value* |
| --- | --- | --- | --- | --- | --- |
|  | 27 (13/14) |  | 27 (13/14) |  |  |
|  | Mean | SD | Mean | SD |  |
| Age | 62.04 | 6.28 | 59.81 | 7.70 | 0.34 |
| ACE-III | 97.89 | 1.80 | 95.41 | 4.21 | 0.06 |
| BDI-II | 4.59 | 4.36 | 15.44 | 11.24 | < 0.001 |
| HADS Dep. | 1.48 | 1.81 | 5.26 | 4.61 | < 0.001 |
| HADS Anx. | 4.30 | 3.16 | 7.04 | 3.32 | < 0.01 |

### Extensive sampling in SCI

As prescribed by rational behaviour, participants in both groups (SCI and healthy controls) adjusted the extent of their search (*s*) to the sampling cost (*η*_*s*_), acquiring fewer samples when this cost increased (Effect of *η*_*s*_ on *s*: *β* = −0.11, 95%*CI* = (−0.14, −0.077), *t*_3232_ = 6.52, *p* < 0.0001, Figure 2a., Table S1). While there was no significant main effect of initial credit (*R*_0_), its interaction with sampling cost was significant (*β* = 0.02, 95%*CI* = (−0.04, −0.004), *t*_3232_ = 2.41, *p* = 0.016, Table S1), which means that the aversive effect of sampling cost on the number of samples obtained was blunted when participants started their search with a larger credit reserve (Figure 2a.).

**Figure 2:**
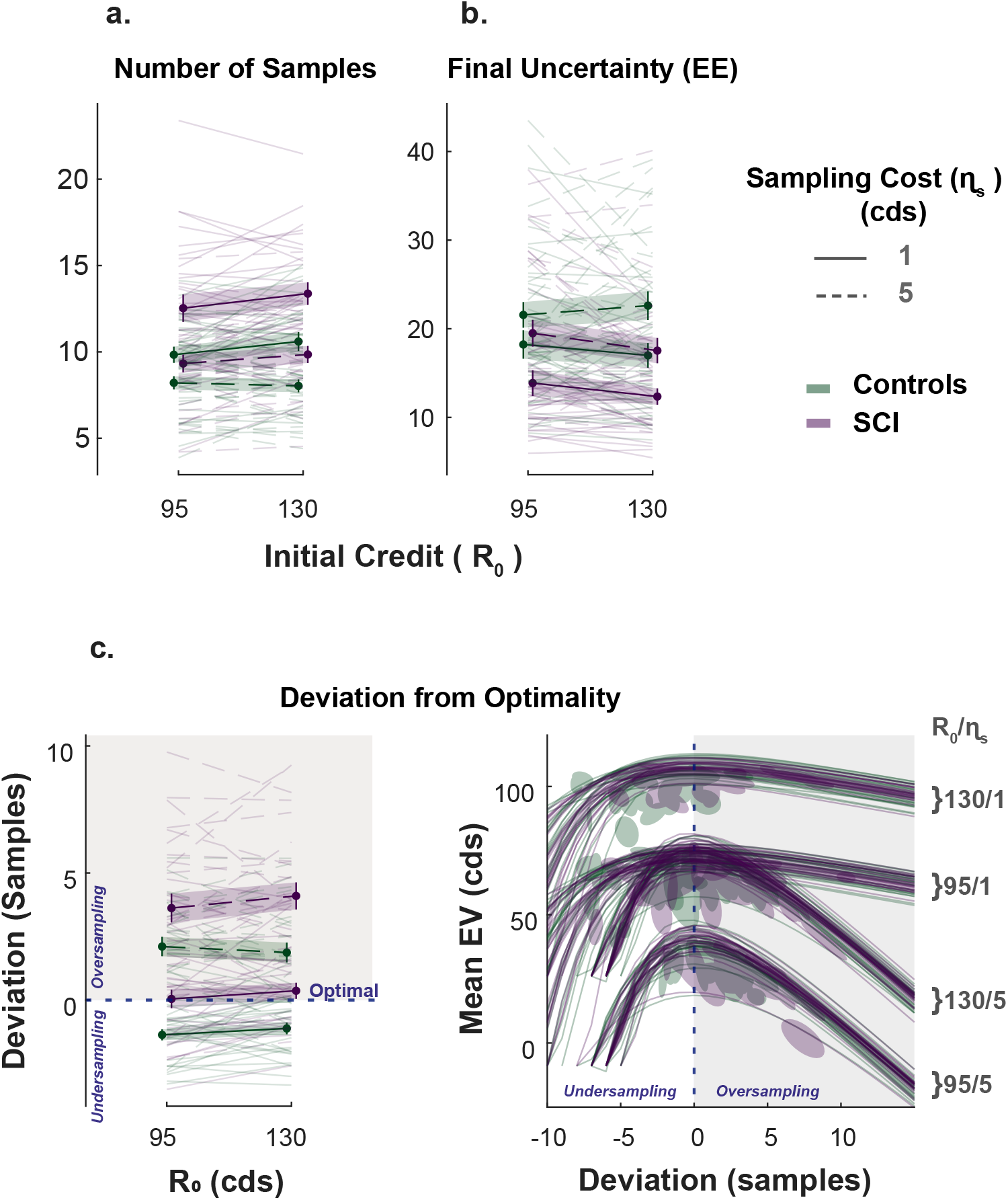
Extensive sampling in SCI. **a.** Across different conditions, individuals with SCI sampled more than healthy controls. **b.** Consequently, SCI participants reached final uncertainty levels (*EE*) lower than controls prior to committing to decisions. **c.** Healthy controls and individuals with SCI over-sampled when sampling cost was high. Over-sampling was more significant in SCI than healthy controls. When sampling cost was low, healthy controls under-sampled while SCI participants were optimal. The panel on the bottom right depicts the changes in expected value (*EV*) as a function of the number of samples deviating from optimal. The optimal number of samples is when *EV* is maximum. Error bars show ±SEM. See Tables S1, S2 & S3 for full statistical details.

The influence of these economic features (*η*_*s*_ and *R*_0_) on the number of samples acquired was not significantly different between SCI participants and controls (*SCI* × *η*_*s*_: *p* = 0.13; *SCI* × *R*_0_: *p* = 0.27). Nevertheless, overall, individuals with SCI sampled significantly more than controls (Main effect of SCI on *s*: *β* = +0.19, 95%*CI* = (0.058, 0.32), *t*_3232_ = 2.81, *p* < 0.01, Figure 2a., Table S1). Gathering more samples led SCI participants to finish their active search at lower levels of uncertainty (*EE*) than controls on average (Main effect of SCI on the *EE* reached at the end of the sampling phase: *β* = 0.26, 95%*CI* = (−0.453, −0.06), *t*_3232_ = 2.66, *p* < 0.01, Table S1), which translated into smaller localisation errors (Main effect of SCI on localisation error: *β* = −0.18, 95%*CI* = (−0.35, −0.004), *t*_3232_ = −2.01, *p* = 0.045).

Next, we asked whether SCI participants’ more extended information gathering led to better performance. To answer this question, we calculated, on each trial, the optimal number of samples, *s*\*, that maximises the expected value of the trial (*EV*). Both acquiring extra samples beyond this point (i.e., over-sampling) and not sampling enough to reach this point (i.e. under-sampling) result in a smaller expected value. Thus, this analysis gave us some insight into the usefulness of the extensive sampling behaviour SCI participants exhibited compared to controls.

Both healthy controls and individuals with SCI over-sampled relative to the optimal stopping point when the sampling cost was high (*p* < 0.001 for both groups, see Table S3 for statistical details). Consistent with above, over-sampling in these conditions was significantly more pronounced in SCI participants compared to controls (Group difference in (*s* − *s\**) at high *η*_*s*_; Low *R*_0_: *t*(52) = 2.066, *p* = 0.04, High *R*_0_: *t*(52) = 3.32, *p* < 0.01). Thus, SCI participants’ tendency to gather more information than controls in these conditions led them to acquire samples with a price outweighing their instrumental benefit.

By contrast, when the sampling cost was low, controls under-sampled relative to the optimal solution (*p* < 0.001 for the two conditions with low *η*_*s*_, see Table S3). Thus, because they acquired more samples in these conditions too, SCI participants better approached optimal sampling behaviour (Group difference in (*s* − *s\**) at low *η*_*s*_; Low *R*_0_: *t*(52) = 3.29, *p* < 0.01, High *R*_0_: *t*(52) = 3.7, *p* < 0.001; Figure 2c.).

To summarise, individuals with SCI sampled more than controls across different experimental conditions, regardless of economic constraints. This was sub-optimal when sampling was expensive (i.e., they overpaid for information) but incidentally led to more optimal behaviour when sampling was cheap.

### Intact passive decision making in individuals with SCI

A passive version of the paradigm was used to investigate what drove SCI participants’ extensive sampling behaviour. More specifically, we tested two hypotheses. First, SCI individuals might have inflated subjective estimates of uncertainty. If this were the case, they might need to reduce uncertainty to a greater extent in order to reach comparable subjective uncertainty levels. Second, SCI participants might have intact estimation of uncertainty but nonetheless assign greater weight to it when balancing it against reward. To test these hypotheses, SCI participants and healthy controls performed a modified version of the paradigm in which they were required to first, report their estimations of experimentally defined levels of uncertainty and second, to accept/reject offers based on whether reward on offer is worth the risk imposed by uncertainty (Figure 3a.).

**Figure 3:**
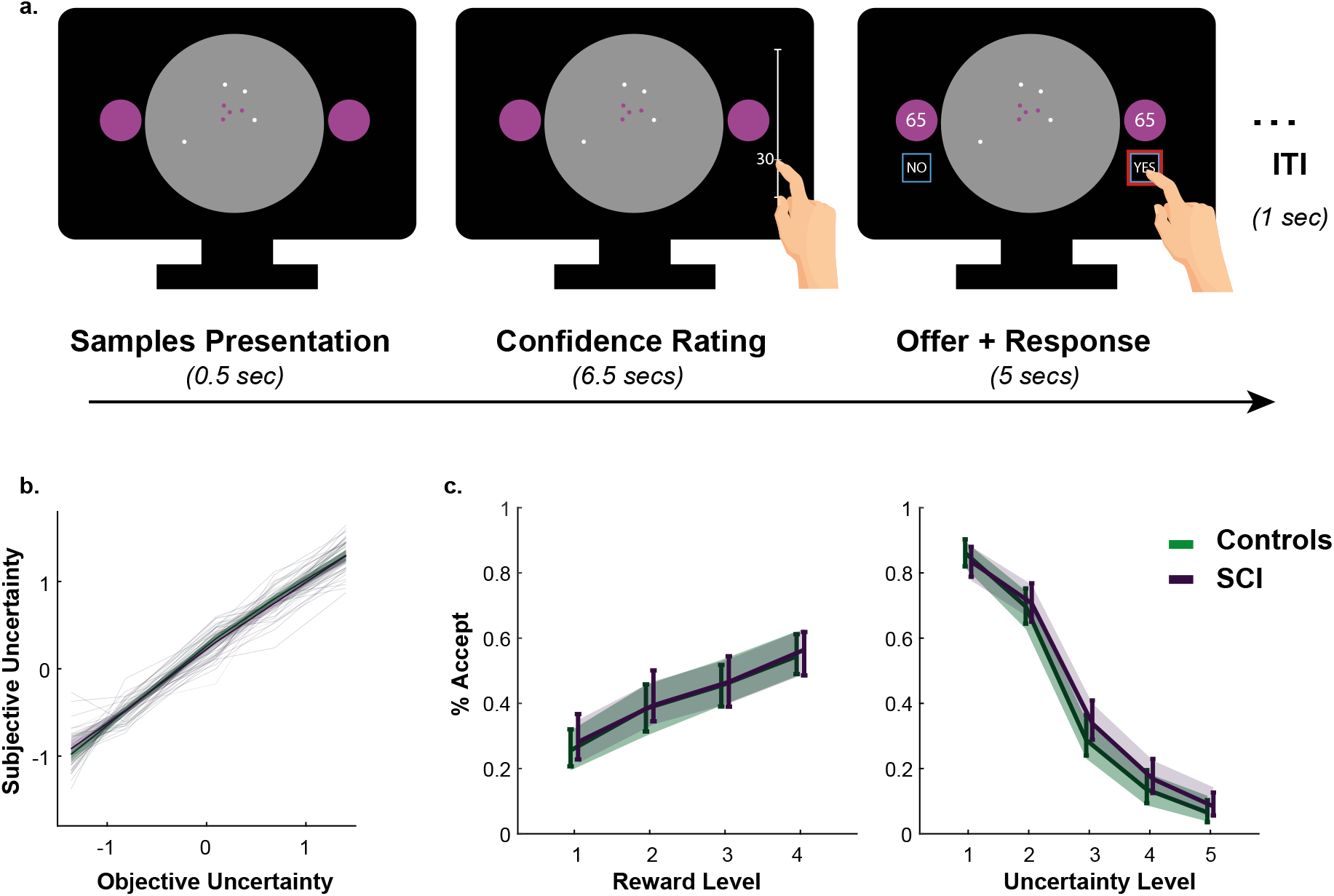
Intact metacognitive judgement and passive decision making in SCI.. **a.** Subjective estimates of uncertainty (z-scored signed-flipped confidence ratings) mapped well onto experimentally defined uncertainty across study participants. There was no significant difference between SCI participants and controls in estimating uncertainty. **b.** There was no significant difference in offer acceptance between individuals with SCI and controls, indicating similar weights assigned to uncertainty and reward when making decisions. Error bars show 95% CI. See Tables S5 & S4 for statistical details.

Generalised mixed effects model was used to investigate the differences in subjective uncertainty between SCI participants and controls. This showed no significant difference in this measure between the two groups (Interaction Group × *EE*: *β* = +0.001, 95%*CI* = (−0.12, 0.11), *t*_5396_ = −0.03, *p* = 0.98, Figure 3b., Table S4), suggesting that tendency to sample more in the active experiment was unlikely caused by biased subjective estimates of uncertainty (i.e., there is no difference in the perceived informational utility of the samples). Similarly, there was no significant difference between the two groups in offer acceptance or in the effects of uncertainty and reward on offer acceptance (Main effect of SCI on offer acceptance: *β* = +0.029, 95%*CI* = (−0.75, 0.80), *t*_5392_ = +0.07, = 0.94; SCI interaction with reward and uncertainty: *p* = 0.70 & *p* = 0.60, respectively, Figure 3c., Table S5). This is consistent with the finding that SCI participants’ extensive sampling behaviour in the active paradigm was mostly independent from economic constraints (no significant interaction SCI ×*R*_0_ or SCI ×*η*_*s*_; Figure 2).

Taken together, these results indicate that extensive sampling in SCI is not related to the way individuals estimate or value uncertainty. Instead, it is likely to capture an intrinsic drive to gather information specifically when agents have agency over the level of uncertainty (i.e. during active sampling) (see *Supplementary information* for a computational model capturing this effect, Figure S1).

### Faster and more efficient sampling in SCI

The key advantage of our paradigm is the possibility to investigate not only how much information people gather but also how quickly and efficiently they do so (Petitet et al., 2021). To capture these extra dimensions of sampling behaviour, we used two behavioural measures: (1) inter-sampling interval, *ISI*, which is the average time interval between consecutive screen touches (shorter *ISI* indicates faster sampling); (2) information extraction rate, *α*, which provides an estimate of the rate at which the *EE* decays over successive samples (higher *α* values indicate higher sampling efficiency).

In young healthy adults, we previously reported a speed-efficiency trade-off whereby slower sampling was associated with greater information extraction rate (i.e., greater reduction of uncertainty at each step of the search) (Petitet et al., 2021). This finding was replicated in the present study (Effect of *ISI* on *α* – Controls: *β* = 0.054, 95%*CI* = (0.032, 0.075), *t*_1618_ = 4.97, *p* < 0.0001; SCI: *β* = 0.052, 95%*CI* = (0.028, 0.076), *t*_1618_ = 4.97, *p* < 0.0001, Figure 4c., Table S8). Investigat-ing group effect using LMM showed that, overall, SCI participants sampled significantly faster than healthy controls (Main effect of group on *ISI*: *β* = −0.29, 95%*CI* = (−0.43, −0.16), *t*_3232_ = −4.24, *p* < 0.0001, Figure 4a., Table S6). Remarkably, despite this faster sampling, SCI participants reduced uncertainty as efficiently as controls (Main effect of group on *α*: *β* = 0.007, 95%*CI* = (−0.015, 0.03), *t*_3232_ = 0.61, *p* = 0.54, Figure 4b., Table S6). In other words, individuals with SCI exceeded the speed-efficiency trade-off that characterised healthy controls’ sampling (Figure 4c.).

**Figure 4:**
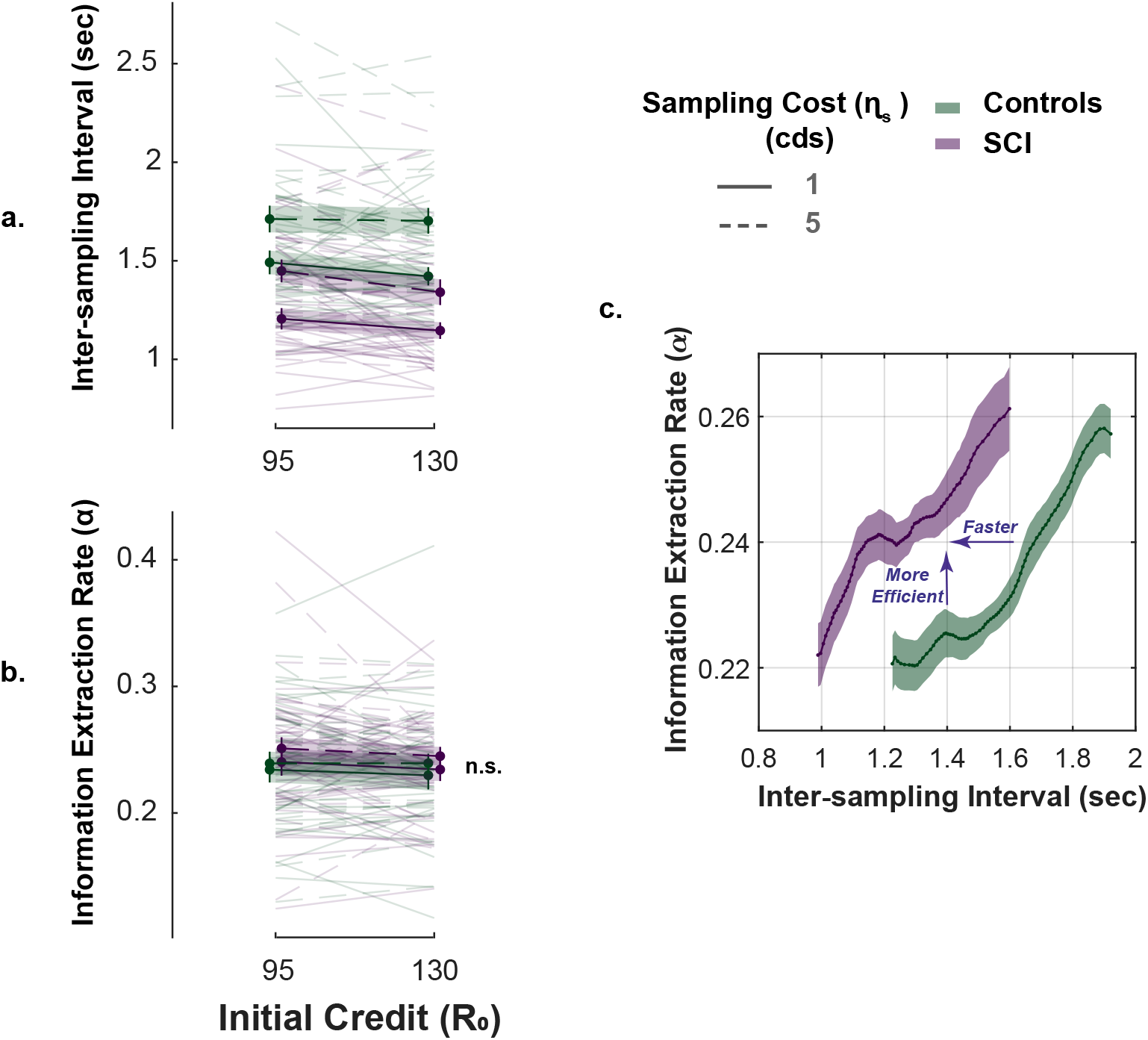
Faster but efficient sampling in SCI. **a.** Across different conditions of the task, SCI participants sampled faster than the healthy controls. **b.** Sampling efficiency was not different between individuals with SCI and control. **c.** Faster sampling was associated with lower efficiency giving rise to a speed-efficiency trade-off. SCI participants exceeded the speed efficiency trade-off that characterised controls’ sampling behaviour as they extracted more information than the control per unit time (sec). Error bars show ±SEM. See Table S6 for full and statistical details.

**Figure 5:**
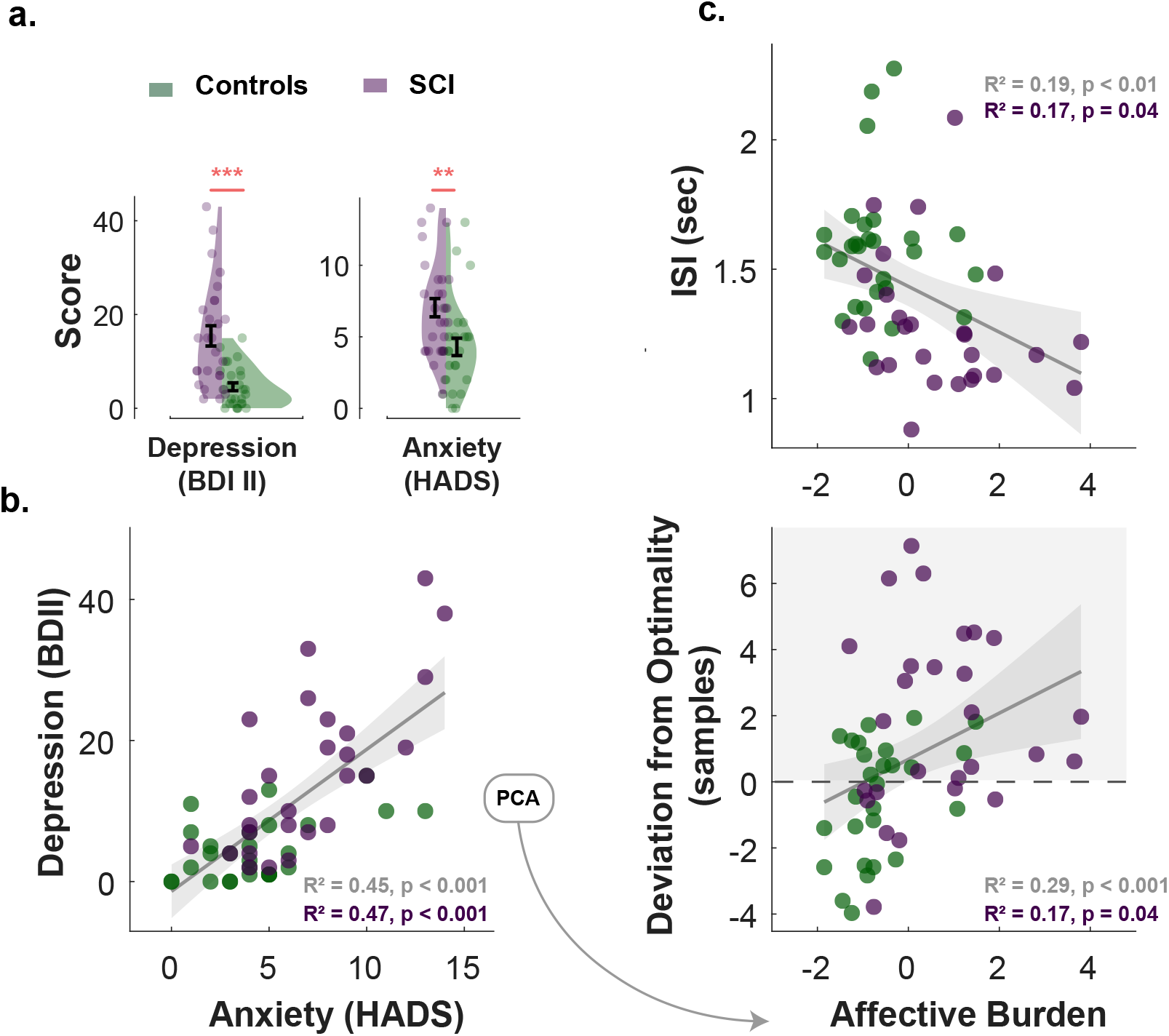
Affective burden correlates with faster and extensive sampling.. **a.** Individuals with SCI were significantly more depressed and anxious than healthy matched control. **b.** Anxiety scores and depression scores significantly correlated with each other across study participants and within SCI group. An affective burden score corresponding to severity of depression and anxiety was extracted using principal component analysis (PCA). This dimension accounted for 84% of the variance between anxiety and depression. **c.** Affective burden was associated with increased sensitivity to uncertainty indexed by speed and extent of sampling. The higher the affective burden score (i.e., the more depressed and anxious a participant is) the faster the sampling rate (shorter *ISI*) and the extensive the sampling was (i.e., over-sampling). These correlations were also significant within SCI group in isolation. BDI II: Beck’s Depression Inventory. HADS: Hospital Anxiety and Depression Scale (only anxiety score was included). ISI: Inter-sampling Interval in seconds. ** : *p* < 0.01, * * * : *p* < 0.001. Error bars show ±SEM. Shaded area in correlation plots show 95% CI

Overall, the performance of individuals with SCI indicates *hypersensitivity to uncertainty*, manifested as more extended, faster though equally efficient information sampling compared to controls.

### Affective burden is associated with more extensive and faster active sampling

Next, we investigated whether markers of hyper-reactivity to uncertainty in SCI (faster and extensive sampling) were associated with affective burden. Non-parametric Spearman’s partial correlations controlled for age and cognitive score across study participants showed that affective burden correlated significantly with sampling speed as well as with deviation from optimal sampling, indicating that both faster and extensive sampling were associated with higher affective burden (Correlation between affective burden and *ISI*: *R*^2^ = 0.19, *p* < 0.01; correlation between affective burden and *s* − *s\**: *R*^2^ = 0.29, *p* < 0.001, Figure 5c.). These correlations were also significant within SCI group in isolation (*R*^2^ = 0.17, *p* = 0.04, for both correlations) as well as within controls group for *s* − *s\** (*R*^2^ = 0.18, *p* = 0.03) but not for ISI (*p* > 0.05).

This result indicates that hyper-reactivity to uncertainty (indexed by faster and extensive sampling) is associated with affective dysregulation in SCI.

### Increased insular-hippocampal connectivity in SCI

Resting-state functional MRI data were collected in 23 SCI participants and 25 controls. We first investigated differences in functional connectivity between SCI participants and controls using a whole-brain network functional connectivity analysis. This entailed investigating the connections between 40 atlas-defined regions of interest (ROIs) that represent the key nodes in brain networks including silence, default mode, sensorimotor, visual, dorsal attention, frontoparietal, language, and cerebellar networks as well as limbic brain regions including the hippocampus, parahippocampus and amygdala (see *Methods*).

Compared to healthy controls, individuals with SCI showed significantly greater functional connectivity between the insular cortex and hippocampal/parahippocampal regions (Insular-hippocampal: *TFCE* = 64.82, *p*_*FWE*_ < 0.01, Insular-parahippocampal: *TFCE* = 70.02, *p*_*FWE*_ < 0.01; Figure 6a., Table S10). There was no other significant difference in resting functional connectivity between any of the other ROIs included in the analysis. Thus, functional connectivity disruptions in SCI participants appeared to be limited and specific to the insular-hippocampal network highlighted in Figure 6a.

**Figure 6:**
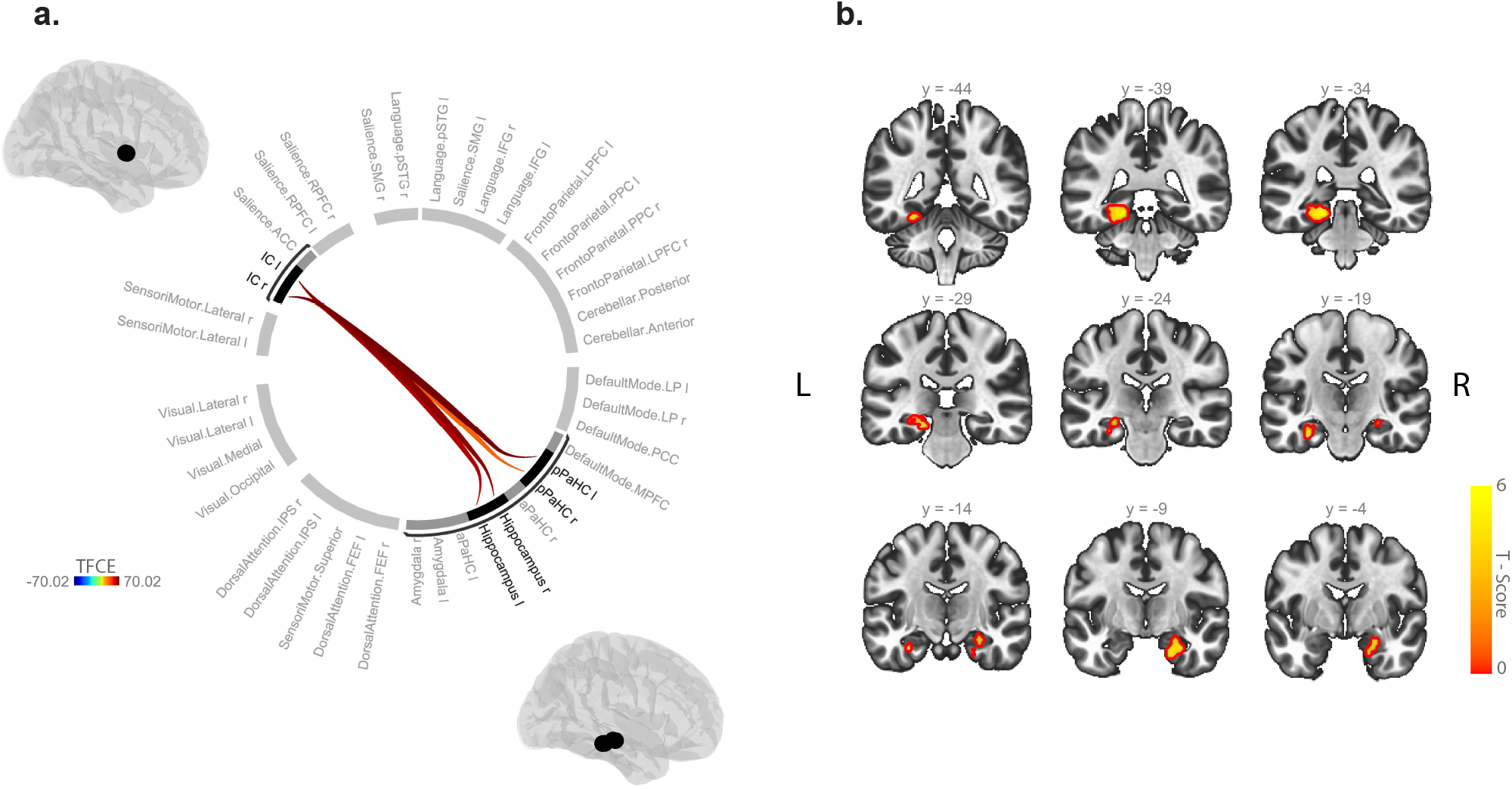
Increased insular hippocampal connectivity in individuals with SCI compared to healthy controls. **a.** Whole-brain ROI-to-ROI functional connectivity analysis with 40 ROIs from different brain networks and regions. SCI participants have increased functional connectivity between insular cortex (IC) and hippocampal/parahippocampal (PaHC) regions. **b.** Seed-based functional connectivity analysis using bilateral insular seed shows voxels in the hippocampal formation with increased functional connectivity with IC in comparison to controls. Analyses were controlled for age and gender. TFCE: Threshold Free Cluster Enhancement. See Tables S9 & S10 for further statistical details.

To complement this analysis, a voxel-wise seed-based functional connectivity analysis was performed using a bilateral insular cortex seed. This aimed to define, at the voxel level, differences in insular functional connectivity between SCI participants and controls (for visualisation purposes). Consistent with the ROI-based analysis, we found that individuals with SCI showed increased functional connectivity between the insular seed and limbic brain regions including hippocampus/parahippocampus and amygdala. Two main significant clusters were found: the first cluster included the left hippocampus (36%) and parahippocampus (38%) (cluster one: *size* = 366 voxels, *p* < 0.001, voxel threshold: *p* < 0.001). The second cluster included the right hippocampus (51%), parahippocampus (26%) and amygdala (20%) (cluster two: *size* = 181 voxels, *p* < 0.01, voxel threshold: *p* < 0.001) (Figure 6b., Table S9).

### Insular-hippocampal connectivity is associated with faster sampling and affective burden

We next investigated the relationship between the strength of insular-hippocampal connectivity and behavioural markers of sensitivity to uncertainty (*ISI* & *s* − *s\**). Non-parametric partial correlations controlling for age, gender, and cognitive score were performed to examine these correlations. The strength of connectivity from ROI-ROI analysis between bilateral insular cortex and hippocampus (right and left) was extracted for this purpose. Multiple correlation testing was corrected using Bonferroni method. The results showed that, across individuals from both groups, stronger bilateral insular connectivity with the right hippocampus significantly correlated with faster sampling rate (*R*^2^ = 0.33, *p* < 0.001, Figure 7) as well as with signed deviation from optimal number of samples (*R*^2^ = 0.22, *p* < 0.001). Within the SCI group, *ISI* but not *s* − *s\** significantly correlated with insular-hippocampal connectivity (Within SCI IC-Hipp connectivity × *ISI*: *R*^2^ = 0.44, *p* < 0.001, × *s* − *s\**: *R*^2^ = 0.09, *p* > 0.05), indicating that sampling speed might be a more sensitive marker of uncertainty sensitivity in this group, despite that these two measures were significantly correlated in this group (*R*^2^ = 0.28, *p* < 0.01).

**Figure 7:**
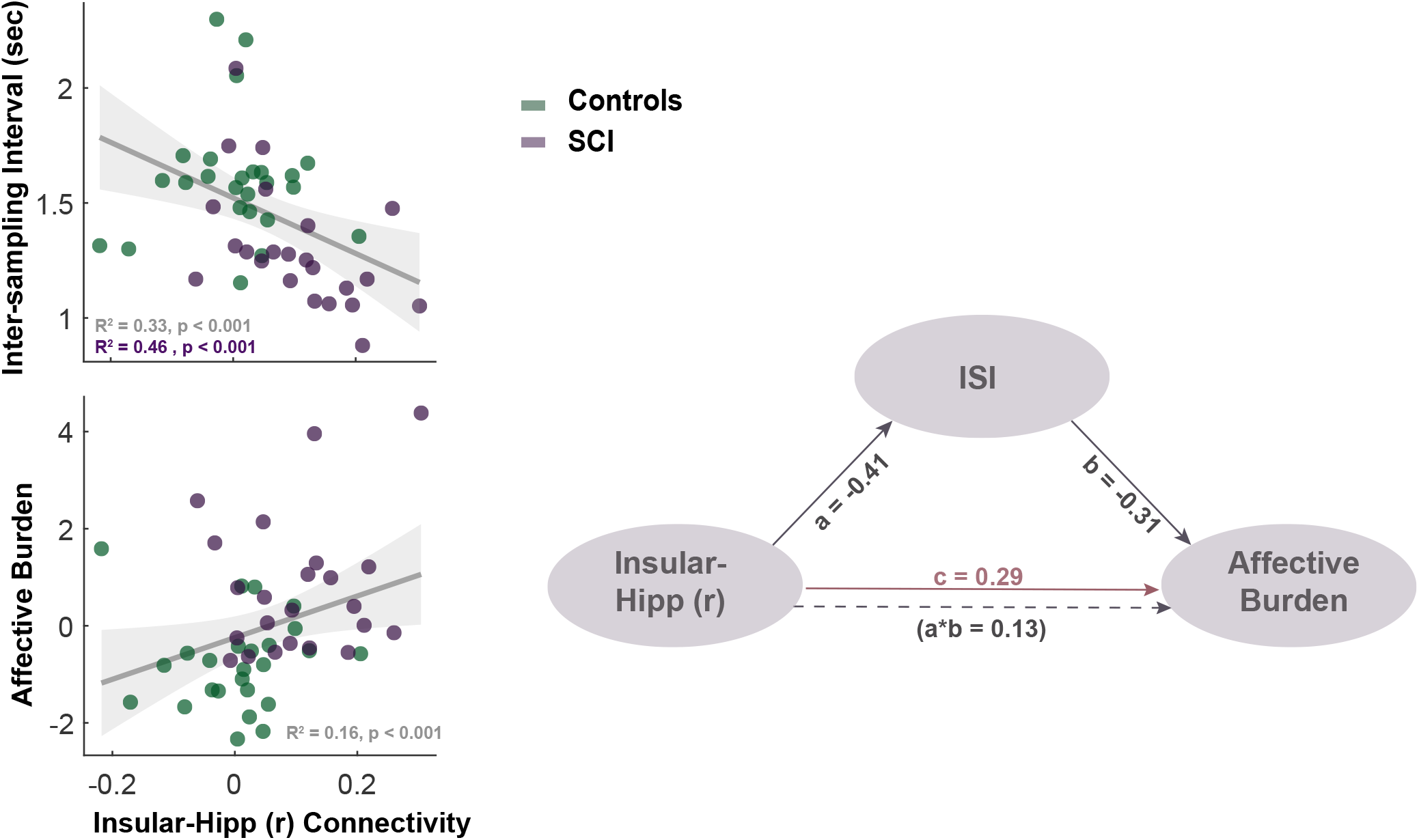
Sensitivity to uncertainty indexed by faster sampling rates mediates the relationship between insular-hippocampal connectivity and affective burden. Across study participants, stronger resting functional connectivity between bilateral insular cortex and right hippocampus was associated with faster sampling rates (i.e., shorter inter-sampling intervals) and higher affective burden. A non-parametric mediation analysis revealed that faster sampling (a behavioural mare of hypersensitivity to uncertainty) mediates the association between insular-hippocampal connectivity and affective burden. Correlations were non-parametric and controlled for age, gender and cognitive score. Shaded area show 95% CI.

Similarly, stronger insular hippocampal connectivity correlated with affective burden across individuals (*R*^2^ = 0.16, *p* < 0.01, Figure 7), however, this correlation within SCI group did not reach significance (*p* > 0.05).

Given that sampling speed correlates with both insular-hippocampal connectivity and affective burden in SCI participants, this indicates that *ISI* as a marker of uncertainty sensitivity might be mediating the correlation between insular-hippocampal connectivity and affective burden. To test this hypothesis we performed a non-parametric causal mediation analysis across study participants (Figure 7). The average causal mediation effect was significant (ACME: *β* = 0.13, 95%*CI* = (0.014, 29), *p* = 0.046, 5000 simulations), indicating that indeed rapid sampling is mediating the relationship between heightened insular-hippocampal connectivity and affective burden across individuals (Prop. Mediated = 0.44). Affective burden score in this analysis was controlled for age, gender, and cognitive score. Caution should be exercised when interpreting these results as these models were performed with data from both groups included. This was done to take advantage of the larger sample size for this type of analysis, while also trying to explore inter-individual differences regardless of group assignment. Such a pooling of data from the two groups is especially feasible with SCI as one operationalisation of the condition considers it part of healthy ageing continuum (Howard, 2020; McWhirter et al., 2020).

## Discussion

A rich body of literature indicates a strong association between SCI and affective dysregulation such as anxiety and depression (Hill et al., 2016; Hohman et al., 2011; Pavisic et al., 2021; Reid and MacLullich, 2006). However, the mechanisms underlying such a burden are not fully established. In this study, we hypothesised that affective dysfunction in SCI might be related to deficits in processing uncertainty. Using a purpose-designed behavioural paradigm, we investigated how people decide and act under un-certainty in active and passive contexts. In the active form, participants could gather information at a cost to reduce uncertainty before committing to decisions. In the passive form, they made decisions responding to offers that had fixed levels of uncertainty and potential reward. The results showed that when participants had agency (i.e., in the active form), individuals with SCI exhibited behaviour indicative of increased reactivity to uncertainty, manifested as more rapid and extensive sampling compared to age- and gender-matched healthy controls. These behavioural markers of heightened reactivity to uncertainty – especially faster sampling rates – correlated with the severity of affective burden in SCI participants. Furthermore, resting functional neuroimaging analysis showed that individuals with SCI had increased insular-hippocampal connectivity, which in turn correlated with uncertainty sensitivity and affective burden across participants. By contrast, estimation and weighing of uncertainty in passive decisions were both intact in SCI participants, indicating that their hypersensitivity to uncertainty was specifically expressed in an active situation in which uncertainty is controllable. Overall, the results point to a specific deficit in processing uncertainty in individuals with SCI that might be underlying their affective dysregulation and is related to increased insular-hippocampal connectivity in the condition.

These results resonate with previous research indicating that people with depression and anxiety might have altered uncertainty processing (Bishop and Gagne, 2018; Boswell et al., 2013; Carleton et al., 2012; Grupe and Nitschke, 2013; Gu et al., 2020; Hartley and Phelps, 2012; Pulcu and Browning, 2019; Saulnier et al., 2019). A recent investigation, for example, showed that a common factor accounting for shared variance between both syndromes was associated with disrupted learning in probabilistic environments reflecting impaired uncertainty processing (Gagne et al., 2020). Other reports pointed to a possible alteration of uncertainty estimation and reward-related valuation in these syndromes affecting decision making under uncertainty (Pizzagalli et al., 2005, 2008; Pulcu and Browning, 2019).

However, evidence on how affective dysfunction relates to more active forms of behaviour such as information gathering prior to committing decisions is limited. One early investigation in social psychology showed that individuals suffering from depression tend to acquire more high-utility information than non-depressed individuals in a simulated interview environment where participants played the role of the interviewer and had to select interview questions from a standardised list of questions that differed in their diagnostic utility (Hildebrand-Saints and Weary, 1989). While this investigation along with other similar reports from social psychology literature advance the notion that disrupted information gathering might be a key feature in affective disorders, a mechanistic account supporting this claim is still not established (Aderka et al., 2013; Camp, 1986; Joiner et al., 1999; Locander and Hermann, 1979). For example, previous studies using classical behavioural paradigms of information seeking such as the beads task failed to report consistent effects of anxiety and depression on performance (Jacoby et al., 2014). Such inconsistencies might be due to the fact that the behavioural paradigms used in these prior studies often neglect an important aspect of information gathering –controllability– which has been hypothesised to be a crucial feature of both anxiety and depression (Abramson et al., 1978; Barlow, 1991). In the present study, this important component was accounted for by using a novel behavioural paradigm allowing participants to gather information with minimal limitations to the speed and efficiency of sampling. This in turn revealed an insightful aspect of information gathering behaviour in individuals with SCI who not only sampled more than controls, but also showed that they do this more rapidly without losing efficiency.

One might argue that rather than suggesting that SCI individuals are hyper-sensitive to uncertainty in the active task, our findings might alternatively be explained by a lower sensitivity to reward – as participants lost more credits to obtain the extra information. However, some observations in the study suggest that this might not be the case. First, the influence of economic constraints (*R*_0_, *η*_*s*_) on sampling behaviour and speed did not significantly differ between individuals with SCI and controls. If SCI participants were less sensitive to reward, then one would expect economic constraints to affect sampling behaviour in SCI participants to a lesser degree than age- and gender-matched controls. Instead, the group effect (i.e., acquiring more samples in individuals with SCI) was equally manifested in all experimental conditions, regardless of the current cost-benefit structure. Second, acquiring more samples indeed allowed SCI participants to achieve lower uncertainty levels prior to decision, suggesting that these additional samples carried instrumental utility and were not merely reflective of wasteful sampling behaviour driven by insensitivity to reward.

Various brain regions have previously been implicated in anxiety and depression and their mechanistic characterisation as deficits in uncertainty processing including amygdala (Grupe and Nitschke, 2013; Morriss et al., 2019), hippocampus (Gray and McNaughton, 2003; Harrison et al., 2006; Rigoli et al., 2019; Strange et al., 2005; Tobia et al., 2012), and insular cortex (Grupe and Nitschke, 2013; Morriss et al., 2019; Tanovic et al., 2018). Consistent with these reports, we found that individuals with SCI displayed heightened connectivity between these regions (insular-limbic). Conceptually, the insula stands out in the context of SCI not only because of its consistent implications in various forms of uncertainty processing and affective syndromes in health and disease (Morriss et al., 2019; Namkung et al., 2017; Paulus and Stein, 2006; Singer et al., 2009; Tanovic et al., 2018), but also because of its potential role in subjective awareness and interoception (Craig, 2009). The insular cortex receives input from different brain regions carrying interoceptive information about various bodily sensations, such as temperature, heartbeat, bowel distension and more (Namkung et al., 2017; Uddin et al., 2017). Subjective experience of these stimuli has been shown to correlate with insular activity on functional neuroimaging using MRI or positron emission tomography (Craig et al., 2000; Critchley et al., 2004). More recent accounts of insular function extend this contribution of subjective awareness to involve emotional states and higher subjective awareness (Chang et al., 2013; Craig, 2009; Namkung et al., 2017). It is thus not surprising to find insular involvement in a condition that is primarily defined by altered subjective experience. A few studies have demonstrated altered insular task-related activity in SCI, however, without functional characterisation of this involvement on goal-directed behaviour and affective functioning (Cai et al., 2020). By contrast, damage to the insula might impair subjectivity and self-awareness as seen in patients with anosognosia for hemiplegia and other forms of insular injury(Fotopoulou et al., 2010; Karnath et al., 2005; Spinazzola et al., 2008), as well as resulting in dysfunctional emotional awareness (e.g., as seen in fronto-temporal dementia), under-reactivity, lack of self-monitoring and passivity (F et al., 1999; Kleiner et al., 2007; VE et al., 2006). Such findings are opposite to what is observed in individuals with SCI who often report heightened levels of subjective affective dysfunction and were found in this study to be behaviourally more reactive.

This insula-centred formalisation of affective dysfunction, uncertainty processing, and subjectivity might be further supported by taking into consideration hippocampal contribution. One prominent view of hippocampal involvement in goal-directed behaviour suggests that it constitutes a crucial part of a behavioural inhibition system (BIS) that is concerned with processing aversive cues such as uncertainty (Gray and McNaughton, 2003). According to this view, the hippocampus acts as a comparator between one’s expectations and the environment, resulting in behavioural aversion to threatening and negative stimuli. It has therefore been suggested that hyperactivity of the BIS might be an important neurobiological basis of anxiety and affective dysregulation (Gray and McNaughton, 2003). When anticipating decisions and actions under uncertainty, the hippocampus might be encoding possible future states and their associated risks (Addis et al., 2011; Martin et al., 2011; Schacter et al., 2008, 2017; Weiler et al., 2010). However, when such situations are avoidable (as in the active task), aversion might be expressed as a propensity to quickly collect information to avoid facing uncertainty in the future. Hippocampal signals encoding future trajectories and prior experiences from these contexts might be shared with the insula, which in turn process emotional responses resulting in a feeling of anxiety and depression (Chang et al., 2013; Craig, 2009; Paulus and Stein, 2006). Additionally, because of the hippocampus’ well-established mnemonic function, insular-hippocampal processing of uncertainty might be also affecting one’s awareness of their memory performance, giving rise to subjective cognitive complaints characterising SCI. Testing such a hypothesis will require future research with a detailed examination of the nature of subjective complaints, their severity, and association with uncertainty and expression of concerns.

In a similar vein, one major future direction is to investigate how the mechanisms and brain networks uncovered in this study relate to the AD spectrum and the prospective risk of developing dementia. For example, while not needed to make a clinical diagnosis of SCI (Jessen et al., 2014), AD-related biological indicators (e.g., CSF biomarkers and amyloid and tau PET imaging) might provide valuable information on how processing uncertainty and information gathering relates to AD pathology in preclinical population with SCI. This could be further supported by evidence from longitudinal follow-up of individuals with SCI to establish risk factors and outcomes. Another line of research might benefit from adopting a transdiagnostic approach to examine whether and how cognitive mechanisms of information seeking and related affective dysfunction are shared (or distinguished) in different stages of AD and other forms of neurodegeneration. Similarly, examining patients who suffer from anxiety and depression without expression of cognitive complaints would help further delineate the association between affective burden and uncertainty-related behaviours.

In conclusion, the study provides evidence suggesting that hyper-reactivity to uncertainty might be a key manifestation of SCI and is related to heightened functional connectivity between the insula and the hippocampus. These manifestations might be underlie affective burden in the condition.

## Materials and methods

### Participants

Twenty seven individuals with SCI (age: *μ* = 59.81, *SD* = 7.70, 14 females) along with 27 healthy age- and gender-matched controls (age: *μ* = 62.04 ± *SD* = 6.28) were recruited for the study. Sample size was determined based on our previous study testing and validating the behavioural paradigm in healthy young controls (Petitet et al., 2021) as well as similar studies investigating information gathering in patient groups (Clark et al., 2006; Hauser et al., 2017). SCI participants were clinically assessed by trained neurologists (co-authors MH & SM) in the cognitive disorders clinic at John Radcliff Hospital, Oxford. In addition to clinical assessment, the diagnosis of SCI was supported by normal performance on standardised objective cognitive assessment using Addenbrooke’s Cognitive Examination (ACE-III) with cutoff > 87/100 (Bruno and Vignaga, 2019; Elamin et al., 2016; Hsieh et al., 2013) as well as normal clinical MRI scan. This definition is consistent with the criteria proposed in previous key reports (Jessen et al., 2014, 2020), suggesting that SCI diagnosis relies on two key components i) subjective report of cognitive decline and ii) normal performance on standardised objective cognitive tests.

All participants gave written consent to take part in the study and were offered monetary compensation for their participation. The study was approved by the University of Oxford ethics committee (RAS ID: 248379, Ethics Approval Reference: 18/SC/0448). Tables 1 shows demographics of the study groups. All participants completed the behavioural tasks and questionnaires. Neuroimaging data were obtained from 23 SCI participants and 25 healthy controls who were MRI compatible and gave consent to be scanned for research purposes.

### Clinical measures

All participants underwent a cognitive assessment using Addenbrooke’s Cognitive Examination III (ACE III) (Hsieh et al., 2013). They also completed self-report questionnaires of depression and anxiety (Beck Depression Inventory-II, BDI-II (Beck et al., 1996) and Hospital Anxiety Depression Scale, HADS (Zigmond and Snaith, 1983)).

### Procedure

A 17-inch touchscreen PC was used to present the task, which was coded using MATLAB (The MathWorks inc., version 2018b) and Psychtoolbox version 3 (Brainard, 1997; Kleiner et al., 2007). The distance between participants and the screen was (~ 50 cm) allowing them to reach it comfortably using their dominant hand. Task environment was adjusted according to handedness and participants were instructed to use their index finger for all their responses. An experimenter was present at all times in the testing room to answer any questions they might have.

### Experimental paradigm

In this study, we used a shorter version of *Circle Quest*, an active information gathering task that has been previously validated and extensively tested in young healthy people (explained in detail in Petitet et al. (2021)). In this paradigm, participants were required to maximise their reward by trying to localise a hidden circle as precisely as possible. They could infer the location of the circle using clues that they could obtain by touching the screen at different locations on a designated search field (grey circle in Figure 1). Participants could acquire as many samples as they wanted without limitations to when and how these samples were obtained on each trial. There were two types of clues: purple dots if the location was situated inside the hidden circle, and white dots if the location was outside the circle. The sizes of hidden circle and dots were fixed on all trials (circle radius: 130 Px, 5.80% of the search space, dot radius: 4 Px). Two circles of the same size as the hidden circle were always displayed on either side of the screen in order to limit memory requirements of the task. Within these two circles was displayed the credits that participants could potentially win if they managed to localise the hidden circle with no errors. After the information-gathering phase, participants could localise the hidden circle using a blue disc that had the same size as the hidden circle.

These aspects of the task were explained to participants using an interactive tutorial with the help of the experimenter. Following this, they performed a training task to further expose them to the task environment and its scoring rules. During training, participants were presented with different configuration of dots (4 purple dots and 4 white dots) from which they could infer the location of the hidden circle with different levels of uncertainty (e.g., when purple dots are spaced out this indicated a lower level of uncertainty than when they were clumped close together). In this training task, participants were instructed to only move the blue disc to where they thought the hidden circle was located. Uncertainty in the task was experimentally quantified as *expected error* (*EE*) which is equal to the error an optimal agent would obtain if they placed the blue disc at the best possible location. A penalty was introduced representing how far localisation was from the true location of the hidden circle. This penalty was subtracted from credits assigned to each trial that participants could potentially win if their localisation was perfect (i.e., placing the blue disc exactly on top of the hidden circle). The penalty incurred for each error pixel was fixed on all trials and was equal to 1.2 credit/pixel, thus localisation error penalty was equal to the distance between the blue disc and hidden circle centre multiplied by 1.2. Once participants completed training, they were required to complete a task comprehension questionnaire to become eligible to continue with the behavioural task. All participants recruited for this study had no issues with this questionnaire.

### Active sampling task

In this version of the task, participants incurred costs for acquiring information: with each additional sample obtained, they lost credits from an initial credit reserve they started each trial with (i.e., from the potential reward they could win if they managed to perfectly find the location of the hidden circle using the blue disc). There were two levels of sampling cost (*η*_*s*_; low: −1 credit/sample and high: −5 credits/sample) and two levels of initial credit (*R*_*o*_; low: 95 credits, high: 130 credit/sample) giving rise to four experimental blocks (15 trials each) that were counterbalanced between participants. Each trial lasted 18 seconds, during which participants could sample the search field freely at any location of the search field. After 18 seconds, the blue disc appeared automatically and participants had six seconds to move it on top of where they thought the hidden circle was located. They then received feedback indicating the number of credits they won on the trial. This score was calculated as follows:

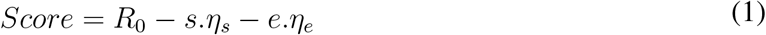

Where *R*_0_ is initial credit reserve, *s* is number of samples acquired, *e* is localisation error (the distance in pixels between the centre of the hidden circle and the centre of the blue disc), and *η*_*e*_ is spatial error cost which was fixed and equal to 1.2 per pixel.

With each additional sample, uncertainty (*EE*) decreased based on the location of the sample and its position in the search trajectory. Efficiency, parameterised as information extraction rate *α*, captures how efficient participants were in reducing *EE* from one sample to the next (see Supplementary Information). Overall, *EE* decreased in the task with successive sampling following an exponential decay function and *α* captures how steep this decline is S3. The expected value, *EV*, at the *s*th sample changed dynamically with acquisition of samples as follows:

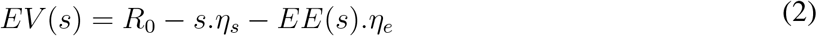

Where *R*_0_ is the initial credit reserve, *s* is the sample number, *η*_*s*_ is the sampling cost, *EE*(*s*) is the expected error (uncertainty) at sample (s), *η*_*e*_ is the spatial error penalty.

Since participants paid a fixed cost for each additional sample (*η*_*s*_) from the initial credit reserve (*R*_0_), this put a limit on the number of samples they could obtain before the information benefit of the sample becomes unjustified by its cost (i.e., imposing a sample number at which the expected value *EV* is maximum). The optimal number of samples *s\** is the number of samples at which *EV* reaches its maximum before it starts declining (i.e., sample benefit become lower than its cost). Deviation from optimal number of samples is simply the difference between the number of samples participants obtain on trial and *s\**. Crucially, this formalisation takes into account differences in sampling efficiency within and between participants, as the utility of the samples acquired is not predefined and depends on where participants choose to touch the screen (i.e., on sampling efficiency). For more details on how these measures are calculated see *Supplementary information* and Petitet et al. (2021)

### Passive choice task

In this version of the task participants’ agency was limited. They were required to make passive decisions (accepting/rejecting offers) based on predetermined levels of uncertainty and reward for these offers. At the beginning of each trial, participants saw a configuration of dots (four purple dots and four white dots) mapping onto different experimentally defined levels of uncertainty (five levels of *EE*: 16.3-24.4, 27.1-38.9, 57.5-58.9, 73.33-74.18, 91.9-93.3 pixels). They were required to indicate how confident they are about the location of the hidden circle using a rating scale on the side of the scene. Subjective uncertainty score was calculated by z-scoring sign-flipping these confidence ratings. Following this, the reward on offer appeared in the two circles on the side of the screen (four reward levels *R*: 40, 65, 90, 115 credits). Participants were required to indicate whether they would like to place the blue disc given the reward and uncertainty of the offers. They did this by pressing ‘Yes’ or ‘No’ appearing on the screen. There were 20 different offer combinations and each offer was presented five times, thus participants completed 100 trials overall. Participants were told that after indicating their preferences, 10 of their accepted offers will be randomly selected for them to play, and that these 10 offers would decide their score in the game.

### Statistical analyses

Statistical analysis was performed either in MATLAB R2019a or R version 4.0.2. Data from active and passive tasks was analysed mainly using generalised mixed effect models with full randomness using *fitglm* function in MATLAB. These were either logistic or linear models depending on the response variable. Full description and statistical details of these models can be found in *Supplementary information*. Post-hoc follow-up analyses were conducted using appropriate statistical tests of difference (student t-test or Wilcoxon rank-sum test) depending on whether parametric assumptions were met. Principal component analysis was applied to HADS anxiety and BDI-II questionnaire scores using *pca* function in MATLAB. The first component of this PCA analysis was used as a measure of affective burden. Correlations with affective burden score were performed using Spearman’s non-parametric testing, controlling for age, gender and objective cognitive score indexed by ACE-III total score. All these correlations were conducted within SCI group and across all individuals included in the study. Mediation analysis testing was performed using *mediation* package(Tingley et al., 2014) in R across all study participants to fulfil sample size needs and characterise behaviour on the spectrum that includes healthy controls and individuals with SCI. Confidence intervals for mediation were estimated using non-parametric bootstrap with bias-corrected and accelerated (BCa) method (5000 simulations) (DiCiccio and Efron, 1996).

### Magnetic Resonance data acquisition

Structural and functional magnetic resonance scans were obtained using 3T Siemens Verio scanner at John Radcliffe Hospital, Oxford. Structural images were T1-weighted with 1 mm isotropic voxel resolution (MPRAGE, field of view: 208 × 256 × 256 matrix, TR/TE = 200/1.94 ms, lip angle = 8°, iPAT =2, prescan-normalise). Resting-state functional MRI (rfMRI) measures spontaneous changes in blood oxygenation (BOLD signal) due to intrinsic brain activity. rfMRI images had voxel size = 2.4 × 2.4 × 2.4 mm (GE-EPI with multi-band acceleration factor = 8, field of view: : 88 × 88 × 64 matrix, TR/TE = 735/39 ms. flip angle = 52°, fat saturation, no iPAT). MR images were obtained from 23/27 SCI participants and 25/27 healthy controls who were MRI compatible or consented to have MR scans as part of the study.

### Magnetic Resonance data processing and analysis

Resting-state connectivity analysis was conducted in MATLAB 2019b using CONN toolbox v20.b (Whitfield-Gabrieli and Nieto-Castanon, 2012) running SPM12. Default processing pipeline was used. This included functional realignment and unwarp, slice-timing and motion correction, segmentation, and normalisation to MNI space. To increase signal-to-noise ratio, spatial smoothing was applied using spatial convolution with Gaussian kernel of 8 mm full width half maximum. Denoising was done using linear regression of potential confounds and temporal band-pass filtering (0.008 - 0.09 Hz). This linear regression controlled for noise components from white matter and cerebrospinal fluid areas (16 parameters each). Head motion was controlled for using 12 noise parameters, which included three translation and three rotation parameters in addition to their first-order derivatives. Confounding effects arising from identified outliers and from linear BOLD signal trends were also controlled for.

Following this, ROI-to-ROI and Seed-to-Voxel analyses were run. For ROI-to-ROI analysis, 40 atlas-defined ROIs were included. These ROIs included the default nodes in CONN used to run network functional connectivity analysis as they represent key nodes in brain networks including silence, default mode, sensorimotor, visual, dorsal attention, frontoparietal, language, and cerebellar networks. In addition, limbic brain regions including the hippocampus, parahippocampus and amygdala were added as these regions have been consistently implicated in processing uncertainty (Harrison et al., 2006; Morriss et al., 2019; Rigoli et al., 2019). Overall, this analysis examined 780 connections, controlling for age and gender differences. Significance testing was done using Threshold Free Cluster Enhancement (TFCE) with corrected connection significance threshold equal to 0.05.

Based on the results of the analysis showing increased insular-hippocampal connectivity in SCI compared to controls (see *Results*), ROI-to-ROI analysis was complemented by seed-to-Voxl analysis seeding from bilateral insular cortex. This aimed to investigate in more detail (at whole-brain voxel level) differences in functional connectivity of the insular cortex between individuals with SCI and matched controls.

## Data and code availability

Anonymised data and code for replicating the main results in the manuscript have been deposited on the Open Science Framework platform: https://osf.io/7ysqu/.

## Supplementary information

### Supplementary methods

#### Quantifying uncertainty and efficiency

Uncertainty in *Circle Quest* paradigm was quantified as expected error (*EE*) which is the average error an ideal participant would obtain upon placing the localisation disc (blue disc) at the best possible location given the dots on the screen (i.e., given the information displayed).

The probability that a location on the screen *λ* is the centre of the hidden circle given the observation *o* at location *σ* was calculated as follows:

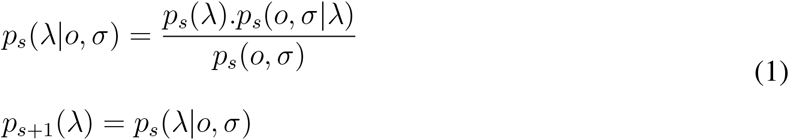

This Bayesian expression is updated sequentially with successive sampling as the posterior probability *p*_*s*_(*λ*|*o, σ*) updates to become the prior probability *p*_*s*+1_(*λ*) with each additional sample.

The likelihood function can be expressed mathematically as:

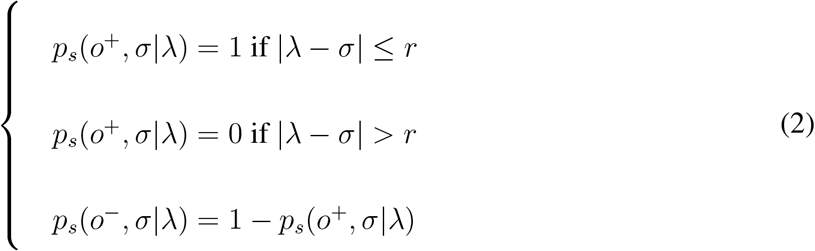

Where *r* is the radius of the hidden circle which is fixed.

Efficiency (information extraction rate, *α*) for each trail per participant was calculated by fitting the following equation:

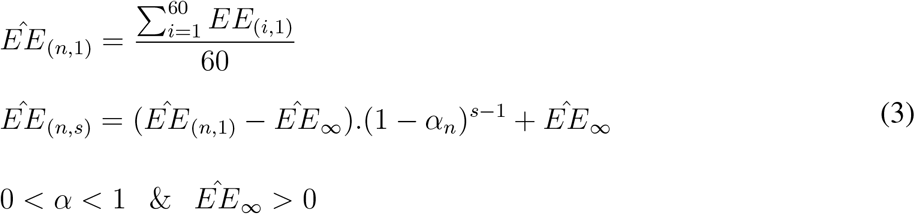

Where *n* is the trial number, *s* is the sample number and 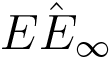 is the asymptotic *EE* which reflects the limitations to uncertainty reduction imposed by the task.

#### Computational modelling of the active task

To further characterise active sampling behaviour, we fitted a previously validated computational model that accounts for hidden cognitive costs in addition to the economic costs imposed by the task when making decisions to sample Petitet et al. (2021). The model calculates the expected utility of a sample (*EU*_*s*_) given these costs and returns four parameter estimates in addition to an intercept per participant. These parameters represent the weights that participants assign to sample cost (*w*_*s*_), sample benefit (*w*_*e*_), speed (*w*_*speed*_), and efficiency (*w*_*α*_) when making decisions to sample.

The model was specified as follows:

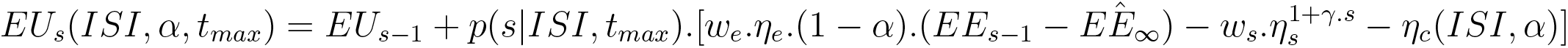

Previous EU + Probability of acquiring the sample given the current time. [Expected information benefit − Sampling cost − Cognitive effort cost]

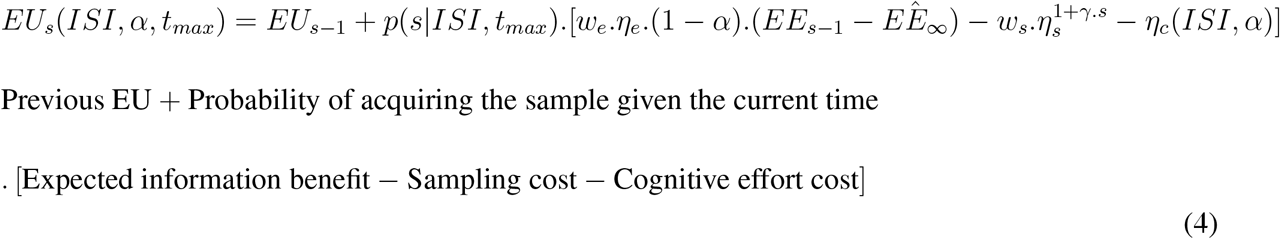

where *γ* is a power term introducing a non-linear transformation of the sampling cost over successive samples and *H*_*c*_(*ISI, α*) is a cognitive effort cost function. 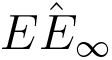 is the information sampling asymptotic limit, which was estimated for each individual beforehand to take into consideration inter-individual variations in asymptotic information sampling performance. For a detailed account of the model specifications and fitting please see Petitet et al. (2021).

Based on previous work (Petitet et al., 2021), a quadratic expression of speed and efficiency was fitted as follows:

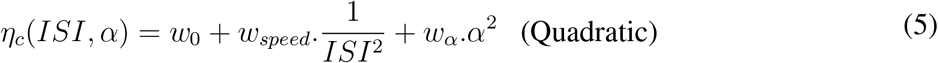

### Supplementary results

#### Individuals with SCI assign lower cognitive costs to both speed and efficiency

The results of the computational model fitted to active search data showed that SCI participants and controls are not significantly different in the weights that they assign sampling cost and benefit (*p >* 0.05 for both, Figure S1), suggesting that extensive sampling in SCI is not driven by task-related economic considerations. Instead, SCI patients, assigned significantly lower costs to sampling speed and efficiency (Group difference – *w*_*speed*_: *z* = −2.35, *p* = 0.018, *w*_*α*_ : = 2.12, *p* = 0.03; Figure S1), indicating that they have lower thresholds to engage in urgent sampling trajectories without losing efficiency.

**Figure S1:**
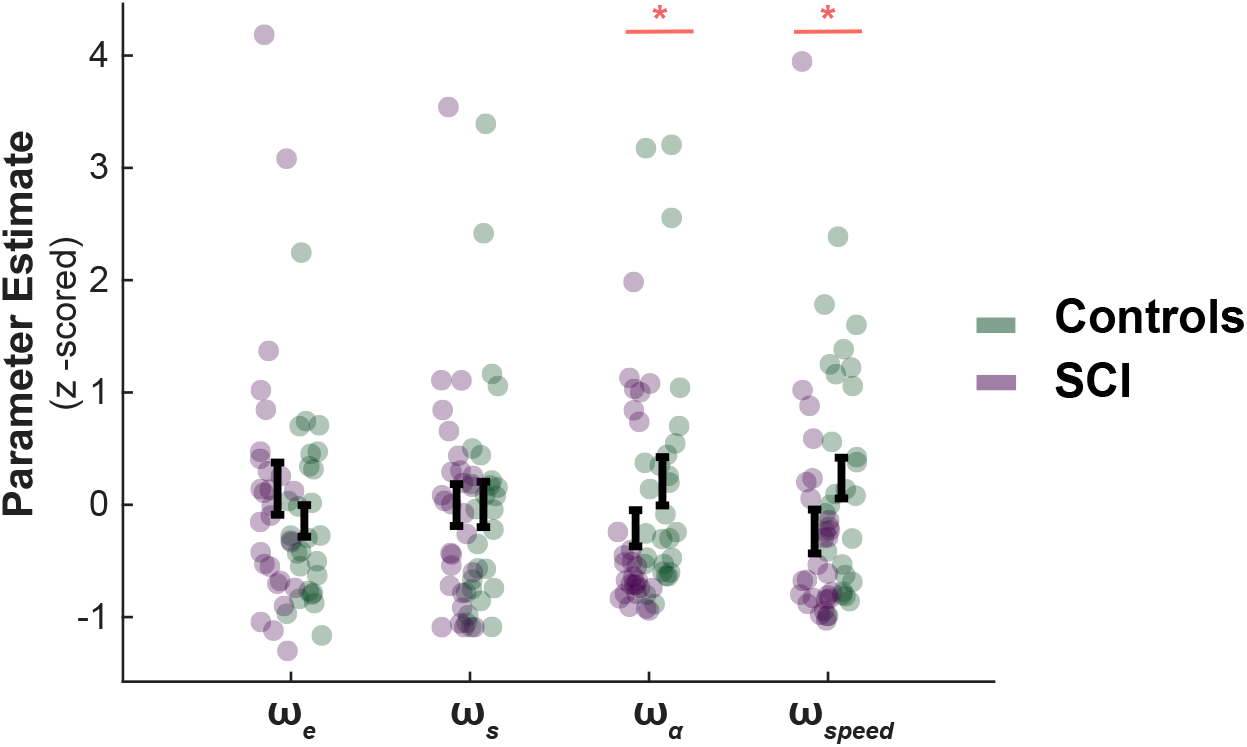
SCI patients assign lower costs to speed and efficiency.. There was no significant difference in the weights SCI participants assigned to sampling benefit (*w*_*e*_) or cost (*w*_*s*_), suggesting the differences in active sampling between the two groups are unlikely to be due to economic constraints of the task. On the other hand, individuals with SCI had lower weights assigned to efficiency (*w*_*α*_) and speed (*w*_*speed*_), indicating a lower cognitive cost to engage in faster and efficient sampling.

**Figure S2:**
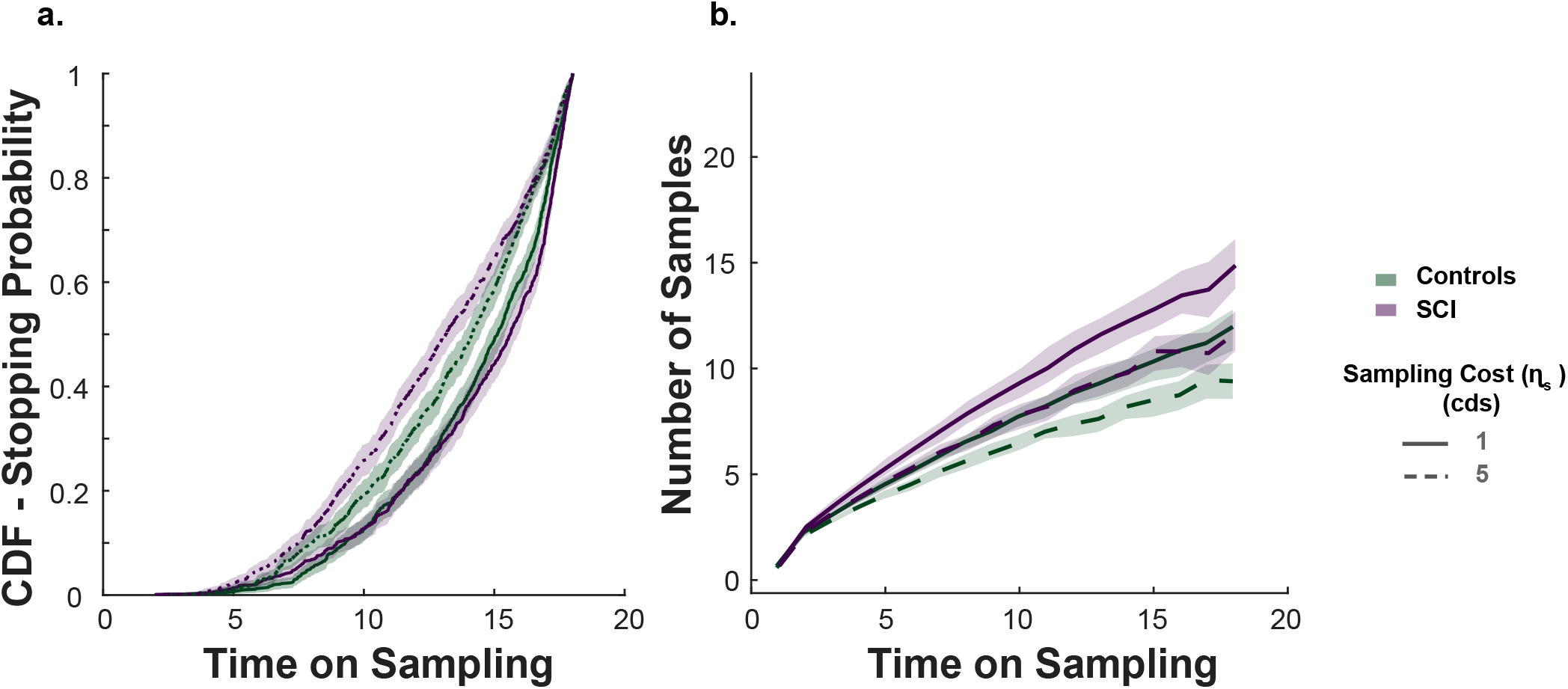
Sampling as a function of time. **a.** Probability to stop sampling at each time point during the sampling phase. When the sampling cost is high, participants are more likely to finish their sampling earlier. This effect was more evident in individuals with SCI as they had faster sampling rates (see Figures 2 & 4). **b.** Number of samples as a function of time. This visualisation shows that SCI participants started to acquire more samples than controls early during the sampling phase. This suggests individuals with SCI opted to decrease uncertainty as soon as possible and that their faster sampling was evident throughout the sampling phase. Shadowed lines show 95% CI.

**Figure S3:**
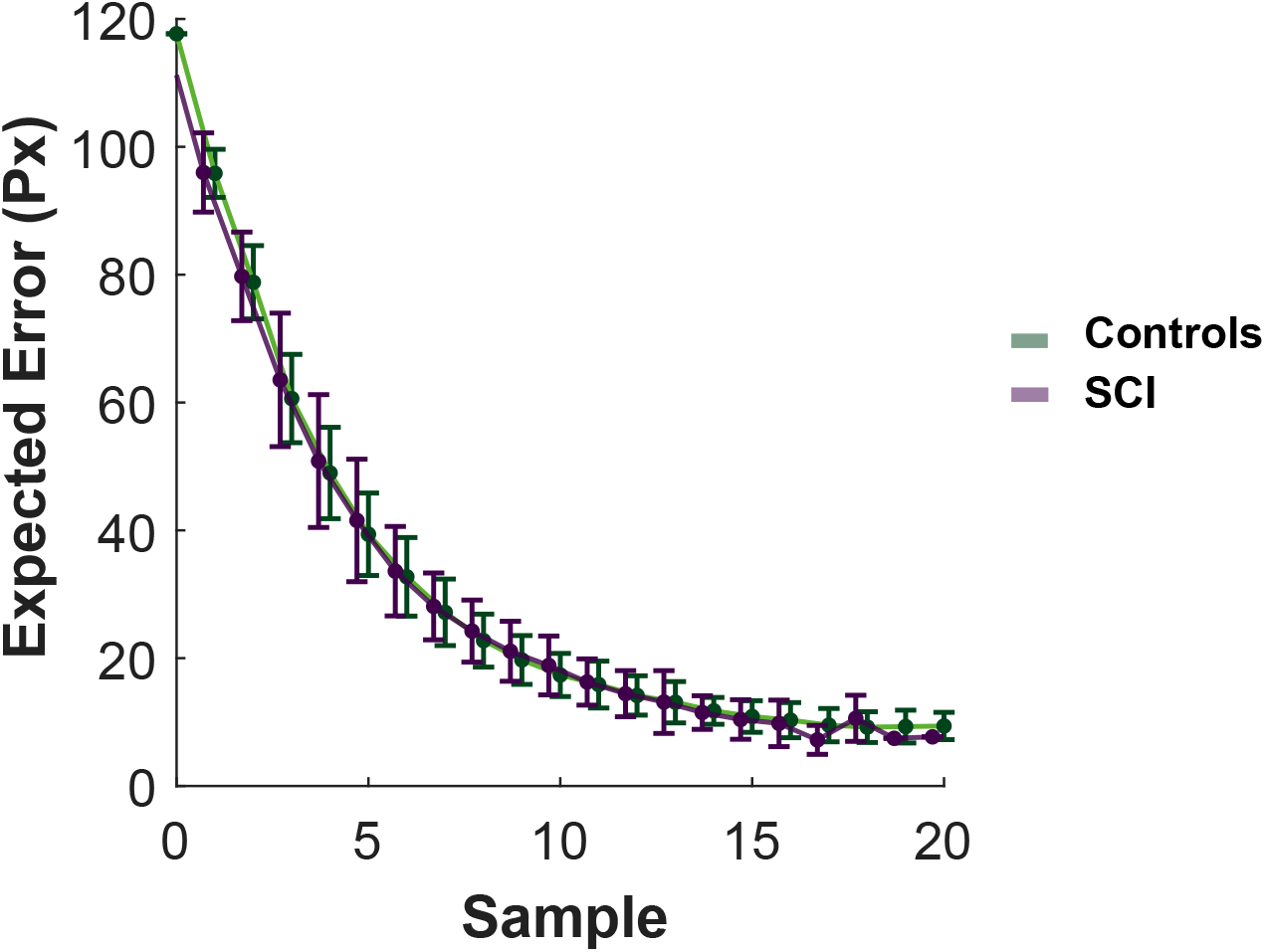
Expected error as a function of sampling. Reduction in uncertainty (expected error, *EE*) with successive sampling follows an exponential decay curve. The samples obtained towards the end of the search have limited benefits compared to the samples obtained earlier. Error bars show ±SEM

**Table S1:** Active Search. Generalised mixed effect models of the effect of the group on performance. Models were specified as follows: Response variable ~ 1 + group\**η*_*s*_ + group\**R*_0_ + *η*_*s*_\**R*_0_ + group:*η*_*s*_:*R*_0_ + (1 + *η*_*s*_\**R*_0_ |participant) SCI: subjective cognitive impairment. *R*_0_: initial credit. *η*_*s*_: sampling cost. *EE*_*end*_: uncertainty before committing to decisions.

| | Samples | $EE_{end}$ | Score | Deviation from Optimal | Expected Value |
| --- | --- | --- | --- | --- | --- |
| (Intercept) | $\beta = +2.18$<br>$SE = 0.0489$<br>$t_{3232} = +44.58$<br>**$p < 0.0001$** | $\beta = +2.83$<br>$SE = 0.0693$<br>$t_{3232} = +40.84$<br>**$p < 0.0001$** | $\beta = +66$<br>$SE = 0.886$<br>$t_{3232} = +74.49$<br>**$p < 0.0001$** | $\beta = -0.48$<br>$SE = 0.432$<br>$t_{3232} = -1.11$<br>$p = 0.27$ | $\beta = +63.3$<br>$SE = 0.697$<br>$t_{3232} = +90.81$<br>**$p < 0.0001$** |
| SCI | $\beta = +0.194$<br>$SE = 0.0691$<br>$t_{3232} = +2.81$<br>**$p = 0.005$** | $\beta = -0.261$<br>$SE = 0.0981$<br>$t_{3232} = -2.66$<br>**$p = 0.0078$** | $\beta = -2.33$<br>$SE = 1.25$<br>$t_{3232} = -1.86$<br>$p = 0.06$ | $\beta = +2.32$<br>$SE = 0.611$<br>$t_{3232} = +3.79$<br>**$p = 0.00015$** | $\beta = -0.197$<br>$SE = 0.985$<br>$t_{3232} = -0.20$<br>$p = 0.84$ |
| SCI:$R_0$ | $\beta = +0.0216$<br>$SE = 0.0196$<br>$t_{3232} = +1.11$<br>$p = 0.27$ | $\beta = -0.0266$<br>$SE = 0.0322$<br>$t_{3232} = -0.83$<br>$p = 0.41$ | $\beta = -0.0693$<br>$SE = 0.618$<br>$t_{3232} = -0.11$<br>$p = 0.91$ | $\beta = +0.203$<br>$SE = 0.206$<br>$t_{3232} = +0.99$<br>$p = 0.32$ | $\beta = +0.0954$<br>$SE = 0.47$<br>$t_{3232} = +0.20$<br>$p = 0.84$ |
| SCI:$\eta_s$ | $\beta = -0.0356$<br>$SE = 0.0237$<br>$t_{3232} = -1.50$<br>$p = 0.13$ | $\beta = +0.0586$<br>$SE = 0.0304$<br>$t_{3232} = +1.93$<br>$p = 0.05$ | $\beta = -1.47$<br>$SE = 1.07$<br>$t_{3232} = -1.37$<br>$p = 0.17$ | $\beta = -0.755$<br>$SE = 0.303$<br>$t_{3232} = -2.49$<br>**$p = 0.013$** | $\beta = -2.86$<br>$SE = 1.01$<br>$t_{3232} = -2.82$<br>**$p = 0.0048$** |
| SCI:$\eta_s$:$R_0$ | $\beta = +0.0189$<br>$SE = 0.0138$<br>$t_{3232} = +1.37$<br>$p = 0.17$ | $\beta = -0.0347$<br>$SE = 0.0231$<br>$t_{3232} = -1.50$<br>$p = 0.13$ | $\beta = +0.261$<br>$SE = 0.678$<br>$t_{3232} = +0.38$<br>$p = 0.70$ | $\beta = +0.128$<br>$SE = 0.155$<br>$t_{3232} = +0.83$<br>$p = 0.41$ | $\beta = -0.0395$<br>$SE = 0.535$<br>$t_{3232} = -0.07$<br>$p = 0.94$ |
| $R_0$ | $\beta = +0.0147$<br>$SE = 0.014$<br>$t_{3232} = +1.05$<br>$p = 0.30$ | $\beta = -0.00661$<br>$SE = 0.0227$<br>$t_{3232} = -0.29$<br>$p = 0.77$ | $\beta = +18.1$<br>$SE = 0.437$<br>$t_{3232} = +41.41$<br>**$p < 0.0001$** | $\beta = +0.0735$<br>$SE = 0.145$<br>$t_{3232} = +0.50$<br>$p = 0.61$ | $\beta = +17.6$<br>$SE = 0.332$<br>$t_{3232} = +52.95$<br>**$p < 0.0001$** |
| $\eta_s$ | $\beta = -0.111$<br>$SE = 0.017$<br>$t_{3232} = -6.52$<br>**$p < 0.0001$** | $\beta = +0.12$<br>$SE = 0.0215$<br>$t_{3232} = +5.60$<br>**$p < 0.0001$** | $\beta = -18.4$<br>$SE = 0.76$<br>$t_{3232} = -24.22$<br>**$p < 0.0001$** | $\beta = +2.25$<br>$SE = 0.214$<br>$t_{3232} = +10.48$<br>**$p < 0.0001$** | $\beta = -17.9$<br>$SE = 0.717$<br>$t_{3232} = -24.92$<br>**$p < 0.0001$** |
| $\eta_s$:$R_0$ | $\beta = -0.0243$<br>$SE = 0.0101$<br>$t_{3232} = -2.41$<br>**$p = 0.016$** | $\beta = +0.0289$<br>$SE = 0.0163$<br>$t_{3232} = +1.77$<br>$p = 0.08$ | $\beta = -0.247$<br>$SE = 0.479$<br>$t_{3232} = -0.52$<br>$p = 0.61$ | $\beta = -0.18$<br>$SE = 0.109$<br>$t_{3232} = -1.64$<br>$p = 0.10$ | $\beta = -0.271$<br>$SE = 0.378$<br>$t_{3232} = -0.72$<br>$p = 0.47$ |
| $adj - R^2$ | 0.77 | 0.44 | 0.72 | 0.82 | 0.89 |
| $N_{obs}$ | 3240 | 3240 | 3240 | 3240 | 3240 |
| AIC | 1.16 | 4887.34 | 27724.93 | 13242.21 | 24174.45 |

**Table S2:** Active Search. Number of samples obtained per condition. *R*_0_: initial credit reserve. *η*_*s*_: sampling cost.

| $R_0/\eta_s$ | Mean | Controls vs. SCI |
| --- | --- | --- |
| 95/1 | $Controls = 9.8346$<br>$SCI = 12.5333$ | $t(52) = -2.9219$<br>$SD = 3.3937$<br>**$p = 0.0051$** |
| 95/5 | $Controls = 8.2074$<br>$SCI = 9.3259$ | $t(52) = -1.7567$<br>$SD = 2.3395$<br>$p = 0.0849$ |
| 130/1 | $Controls = 10.6000$<br>$SCI = 13.3753$ | $t(52) = -3.2560$<br>$SD = 3.1318$<br>**$p = 0.0020$** |
| 130/5 | $Controls = 8.0321$<br>$SCI = 9.8494$ | $t(52) = -2.9422$<br>$SD = 2.2694$<br>**$p = 0.0049$** |

**Table S3:** Active Search. Deviation from optimal (s - *s\** for each condition. *R*_0_: initial credit reserve. *η*_*s*_: sampling cost.

| $R_0/\eta_s$ | Controls | SCI | Controls vs. SCI |
| --- | --- | --- | --- |
| 95/1 | $\beta = -2.98$<br>$SE = 0.47$<br>$t_{1616} = -6.34$<br>**$p < 0.0001$** | $\beta = +0.0148$<br>$SE = 0.757$<br>$t_{1616} = +0.02$<br>$p = 0.98$ | $Mean_{Controls} = -2.9802$<br>$Mean_{SCI} = -2.9802$<br>$t(52) = -3.2984$<br>**$p = 0.0018$** |
| 95/5 | $\beta = +1.87$<br>$SE = 0.334$<br>$t_{1616} = +5.61$<br>**$p < 0.0001$** | $\beta = +3.1$<br>$SE = 0.48$<br>$t_{1616} = +6.47$<br>**$p < 0.0001$** | $Mean_{Controls} = +1.8741$<br>$Mean_{SCI} = +3.1037$<br>$t(52) = -2.0644$<br>**$p = 0.0440$** |
| 130/1 | $\beta = -2.47$<br>$SE = 0.523$<br>$t_{1616} = -4.73$<br>**$p < 0.0001$** | $\beta = +0.672$<br>$SE = 0.627$<br>$t_{1616} = +1.07$<br>$p = 0.28$ | $Mean_{Controls} = -2.4741$<br>$Mean_{SCI} = +0.6716$<br>$t(52) = 3.7820$<br>**$p = 0.0004$** |
| 130/5 | $\beta = +1.66$<br>$SE = 0.337$<br>$t_{1616} = +4.93$<br>**$p < 0.0001$** | $\beta = +3.55$<br>$SE = 0.457$<br>$t_{1616} = +7.77$<br>**$p < 0.0001$** | $Mean_{Controls} = +1.6617$<br>$Mean_{SCI} = +3.5531$<br>$t(52) = -3.2668$<br>**$p = 0.0019$** |
| $adj - R^2$ | 0.83 | 0.77 | |
| $N_{obs}$ | 1620 | 1620 | |
| AIC | 6117.11 | 7013.05 |  |

**Table S4:**
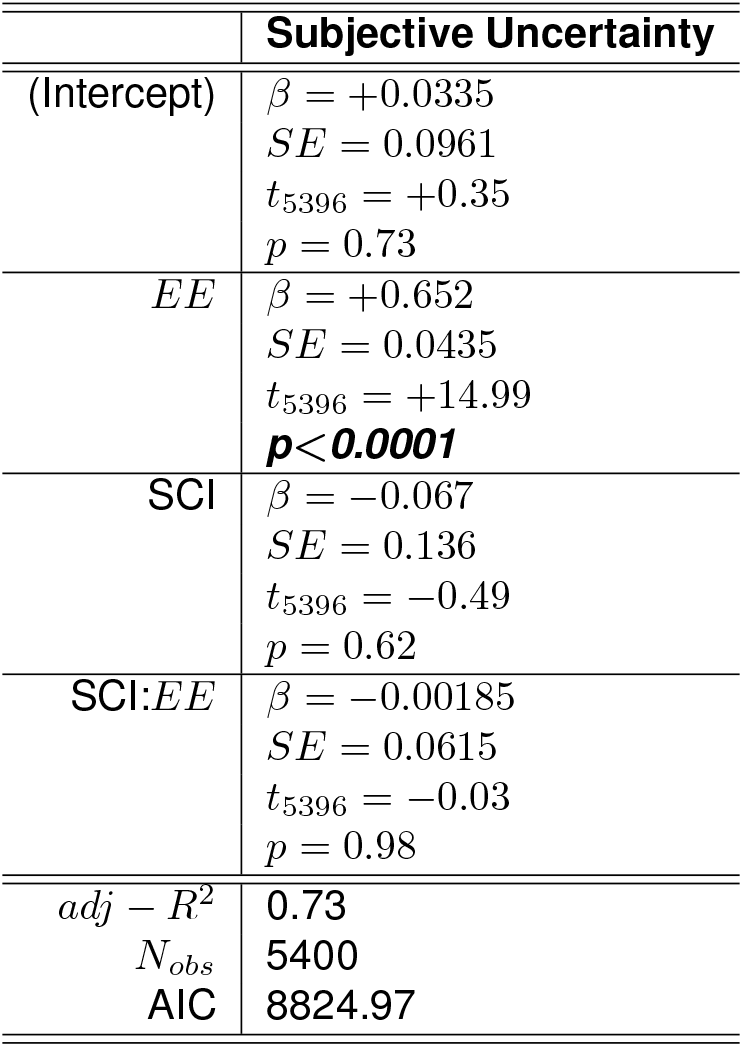
Active Search. Generalised mixed effect models of the effect of the group on accuracy of uncertainty estimation. Models were specified as follows. Subjective Uncertainty: Subjective Uncertainty ~ 1 + group\**EE* + (1 + *EE* |participant) + (1 |trial).

|  | Subjective Uncertainty |
| --- | --- |
| (Intercept) | $\beta = +0.0335$<br>$SE = 0.0961$<br>$t_{5396} = +0.35$<br>$p = 0.73$ |
| EE | $\beta = +0.652$<br>$SE = 0.0435$<br>$t_{5396} = +14.99$<br>**$p < 0.0001$** |
| SCI | $\beta = -0.067$<br>$SE = 0.136$<br>$t_{5396} = -0.49$<br>$p = 0.62$ |
| SCI:EE | $\beta = -0.00185$<br>$SE = 0.0615$<br>$t_{5396} = -0.03$<br>$p = 0.98$ |
| $adj - R^2$ | 0.73 |
| $N_{obs}$ | 5400 |
| AIC | 8824.97 |

**Table S5:** Generalised mixed effect model investigating group difference in passive decision making under uncertainty. Models were specified as follows. Passive Choices: choice ~ 1 + group\**R* + group\**EE* + *R*\**EE* + group:*R*:*EE* + (1 + *R*\**EE* |participant). *R*: reward on offer. *EE*: uncertainty of offer represented by dot configurations and computed as expected localisation error. SCI: subjective cognitive impairment group.

|  | Passive Choices |
| --- | --- |
| (Intercept) | $\beta = -0.416$<br>$SE = 0.281$<br>$t_{5392} = -1.48$<br>$p = 0.14$ |
| SCI | $\beta = +0.029$<br>$SE = 0.397$<br>$t_{5392} = +0.07$<br>$p = 0.94$ |
| SCI:EE | $\beta = +0.167$<br>$SE = 0.318$<br>$t_{5392} = +0.53$<br>$p = 0.60$ |
| SCI:R | $\beta = -0.103$<br>$SE = 0.272$<br>$t_{5392} = -0.38$<br>$p = 0.71$ |
| SCI:R:EE | $\beta = +0.169$<br>$SE = 0.14$<br>$t_{5392} = +1.21$<br>$p = 0.23$ |
| EE | $\beta = -2.77$<br>$SE = 0.225$<br>$t_{5392} = -12.31$<br>**$p < 0.0001$** |
| R | $\beta = +1.12$<br>$SE = 0.193$<br>$t_{5392} = +5.80$<br>**$p < 0.0001$** |
| R:EE | $\beta = -0.0493$<br>$SE = 0.103$<br>$t_{5392} = -0.48$<br>$p = 0.63$ |
| $adj - R^2$ | 0.96 |
| $N_{obs}$ | 5400 |
| AIC | 3959.72 |

**Table S6:** Active Search. Generalised mixed effect models of the effect of the group on sampling speed (ISI) and efficiency (*α*). Models were specified as follows. Inter-sampling Interval: *ISI* ~ 1 + group\**η*_*s*_ + group\**R*_0_ + *η*_*s*_\**R*_0_ + group:*η*_*s*_:*R*_0_ + (1 + *η*_*s*_\**R*_0_ |participant); Information extraction rate: *α* ~ 1 + group\**η*_*s*_ + group\**R*_0_ + *η*_*s*_\**R*_0_ + group:*η*_*s*_:*R*_0_ + (1 + *η*_*s*_\**R*_0_ |participant).

| | *ISI* | $\alpha$ |
| --- | --- | --- |
| (Intercept) | $\beta = +1.58$<br>$SE = 0.0495$<br>$t_{3232} = +31.98$<br>**$p < 0.0001$** | $\beta = +0.236$<br>$SE = 0.00812$<br>$t_{3232} = +29.03$<br>**$p < 0.0001$** |
| SCI | $\beta = -0.297$<br>$SE = 0.07$<br>$t_{3232} = -4.24$<br>**$p < 0.0001$** | $\beta = +0.00702$<br>$SE = 0.0115$<br>$t_{3232} = +0.61$<br>$p = 0.54$ |
| SCI:$R_0$ | $\beta = -0.0221$<br>$SE = 0.0247$<br>$t_{3232} = -0.90$<br>$p = 0.37$ | $\beta = -0.00195$<br>$SE = 0.00372$<br>$t_{3232} = -0.52$<br>$p = 0.60$ |
| SCI:$\eta_s$ | $\beta = -0.0161$<br>$SE = 0.0265$<br>$t_{3232} = -0.61$<br>$p = 0.54$ | $\beta = +0.00172$<br>$SE = 0.00389$<br>$t_{3232} = +0.44$<br>$p = 0.66$ |
| SCI:$\eta_s$:$R_0$ | $\beta = -0.0274$<br>$SE = 0.0165$<br>$t_{3232} = -1.66$<br>$p = 0.10$ | $\beta = -0.00117$<br>$SE = 0.00309$<br>$t_{3232} = -0.38$<br>$p = 0.71$ |
| $R_0$ | $\beta = -0.0199$<br>$SE = 0.0175$<br>$t_{3232} = -1.14$<br>$p = 0.26$ | $\beta = -0.00108$<br>$SE = 0.00263$<br>$t_{3232} = -0.41$<br>$p = 0.68$ |
| $\eta_s$ | $\beta = +0.125$<br>$SE = 0.0187$<br>$t_{3232} = +6.70$<br>**$p < 0.0001$** | $\beta = +0.0036$<br>$SE = 0.00275$<br>$t_{3232} = +1.31$<br>$p = 0.19$ |
| $\eta_s$:$R_0$ | $\beta = +0.0153$<br>$SE = 0.0117$<br>$t_{3232} = +1.31$<br>$p = 0.19$ | $\beta = +0.00113$<br>$SE = 0.00219$<br>$t_{3232} = +0.52$<br>$p = 0.60$ |
| $adj - R^2$ | **0.64** | **0.18** |
| $N_{obs}$ | **3240** | **3240** |
| AIC | **1059.49** | **-6438.06** |

**Table S7:** Active Search – Inter-sampling interval per condition. *R*_0_: initial credit reserve. *η*_*s*_: sampling cost.

| $R_0/\eta_s$ | Mean | Controls vs. SCI |
| --- | --- | --- |
| 95/1 | $Controls = 1.4917$<br>$SCI = 1.2056$ | $z = 3.7195$<br>$ranksum = 958$<br>**$p = 0.0002$** |
| 95/5 | $Controls = 1.7119$<br>$SCI = 1.4485$ | $z = 3.0275$<br>$ranksum = 918$<br>**$p = 0.0025$** |
| 130/1 | $Controls = 1.4214$<br>$SCI = 1.1458$ | $t(52) = 4.4093$<br>$SD = 0.2296$<br>**$p = 0.0001$** |
| 130/5 | $Controls = 1.7028$<br>$SCI = 1.3404$ | $z = 3.8060$<br>$ranksum = 963$<br>**$p = 0.0001$** |

**Table S8:** Active Search. Speed efficiency trade-off. Models were specified as follows. *α* ~ 1 + *ISI* + (1 + *ISI* |participant) + (1 + *ISI* |condition) + (1 |trial).

|  | Controls | SCI |
| --- | --- | --- |
| (Intercept) | $\beta = +0.152$<br>$SE = 0.0168$<br>$t_{1618} = +9.01$<br>**$p < 0.0001$** | $\beta = +0.174$<br>$SE = 0.0173$<br>$t_{1618} = +10.06$<br>**$p < 0.0001$** |
| *ISI* | $\beta = +0.0541$<br>$SE = 0.0109$<br>$t_{1618} = +4.97$<br>**$p < 0.0001$** | $\beta = +0.0524$<br>$SE = 0.0125$<br>$t_{1618} = +4.21$<br>**$p < 0.0001$** |
| $adj - R^2$ | 0.23 | 0.20 |
| $N_{obs}$ | 1620 | 1620 |
| AIC | -3296.91 | -3258.57 |

**Table S9:** Seed-based resting functional connectivity with bilateral insular cortex seed. Table shows significant clusters. FWE: family-wise error. FDR: false discovery rate.

|  | x,y,z | size | size p-FWE | size p-FDR | size p-unc |
| --- | --- | --- | --- | --- | --- |
| **Cluster 1** | -18 -34 -14 | 366 | 0.000003 | 0.000004 | 0.000000 |
| **Cluster 2** | +28 -08 -24 | 181 | 0.001251 | 0.001021 | 0.000035 |

**Table S10:** ROI to ROI resting functional connectivity. PaHC: parahippocampus. Hipp: hippocampus. IC: insular cortex. unc: uncorrected. FWE: family-wise error. FDR: false discovery rate.

| Analysis Unit | Statistic | p-unc | p-FDR | p-FWE |
| --- | --- | --- | --- | --- |
| **Cluster 1** | TFCE = 70.02 | 0.000025 | 0.001730 | 0.002000 |
| Connection pPaHC l - IC r | T(44) = 4.63 | 0.000032 | 0.017440 |  |
| Connection pPaHC l - IC l | T(44) = 4.45 | 0.000058 | 0.017440 |  |
| Connection pPaHC r - IC l | T(44) = 3.09 | 0.003464 | 0.150095 |  |
| **Cluster 2** | TFCE = 64.82 | 0.000029 | 0.001730 | 0.003000 |
| Connection Hipp r - IC l | T(44) = 4.40 | 0.000067 | 0.017440 |  |
| Connection Hipp l - IC r | T(44) = 4.07 | 0.000195 | 0.038090 |  |
| Connection Hipp l - IC l | T(44) = 3.74 | 0.000534 | 0.083315 |  |

## References

Lyn Y. Abramson, Martin E. Seligman, and John D. Teasdale. Learned helplessness in humans: Critique and reformulation. Journal of Abnormal Psychology, 87(1):49–74, 2 1978. ISSN 0021843X. doi: 10.1037/0021-843X.87.1.49. URL /record/1979-00305-001?doi=1.

Donna Rose Addis, Theresa Cheng, Reece P. Roberts, and Daniel L. Schacter. Hippocampal contributions to the episodic simulation of specific and general future events. Hippocampus, 21(10):1045–1052, 10 2011. ISSN 10509631. doi: 10.1002/HIPO.20870. URL https:///pmc/articles/PMC3815461//pmc/articles/PMC3815461/?report=abstracthttps://www.ncbi.nlm.nih.gov/pmc/articles/PMC3815461/.

Idan M. Aderka, Ayala Haker, Sofi Marom, and Haggai Hermesh. Brief report information-seeking bias in social anxiety disorder. Journal of Abnormal Psychology, 122(1):7–12, 2013. ISSN 19391846. doi: 10.1037/a0029555. URL https://pubmed.ncbi.nlm.nih.gov/22905860/.

Kelly Allott, Caroline Gao, Sarah E. Hetrick, Kate M. Filia, Jana M. Menssink, Caroline Fisher, Ian B. Hickie, Helen E. Herrman, Debra J. Rickwood, Alexandra G. Parker, Patrick D. Mcgorry, and Sue M. Cotton. Subjective cognitive functioning in relation to changes in levels of depression and anxiety in youth over 3 months of treatment. BJPsych Open, 6(5), 9 2020. ISSN 2056-4724. doi: 10.1192/bjo.2020. URL /pmc/articles/PMC7453798//pmc/articles/PMC7453798/?report=abstracthttps://www.ncbi.nlm.nih.gov/pmc/articles/PMC7453798/.

Zoe Arvanitakis, Sue E. Leurgans, Debra A. Fleischman, Julie A. Schneider, Kumar B. Rajan, Jeremy J. Pruzin, Raj C. Shah, Denis A. Evans, Lisa L. Barnes, and David A. Bennett. Memory complaints, dementia, and neuropathology in older blacks and whites. Annals of Neurology, 83(4):718–729, 4 2018. ISSN 1531-8249. doi: 10.1002/ANA.25189. URL https://onlinelibrary.wiley.com/doi/full/10.1002/ana.25189 https://onlinelibrary.wiley.com/doi/abs/10.1002/ana.25189https://onlinelibrary.wiley.com/doi/10.1002/ana.25189.

Dominik R. Bach and Raymond J. Dolan. Knowing how much you don’t know: a neural organization of uncertainty estimates. Nature Reviews Neuroscience, 13(8):572–586, 8 2012. ISSN 1471-003X. doi: 10.1038/nrn3289. URL http://www.nature.com/articles/nrn3289.

David Harrison Barlow. Disorders of Emotion. Psychological Inquiry, 2(1):58–71, 1991. ISSN 1532-7965. doi: 10.1207/s15327965pli0201{\}15. URL https://www.tandfonline.com/action/journalInformation?journalCode=hpli20.

Aaron T. Beck, Robert A. Steer, Roberta Ball, and William F. Ranieri. Comparison of Beck depression inventories −IA and −II in psychiatric outpatients. Journal of Personality Assessment, 67(3):588–597, 12 1996. ISSN 00223891. doi: 10.1207/s15327752jpa6703{\}13. URL https://www.tandfonline.com/doi/abs/10.1207/s15327752jpa6703_13.

Sonia J. Bishop and Christopher Gagne. Anxiety, Depression, and Decision Making: A Computational Perspective. Annual Review of Neuroscience, 41(1):371–388, 2018. ISSN 0147-006X. doi: 10.1146/annurev-neuro-080317-062007.

Paul A. Boelen, Albert Reijntjes, and Geert E. Smid. Concurrent and prospective associations of intolerance of uncertainty with symptoms of prolonged grief, posttraumatic stress, and depression after bereavement. Journal of Anxiety Disorders, 41:65–72, 6 2016. ISSN 0887-6185. doi: 10.1016/J.JANXDIS.2016.03.004.

James F Boswell, Johanna Thompson-Hollands, Todd J Farchione, and David H Barlow. Intolerance of Uncertainty: A Common Factor in the Treatment of Emotional Disorders. 2013. doi: 10.1002/jclp.21965.

David H. Brainard. The Psychophysics Toolbox. Spatial Vision, 10(4):433–436, 1 1997. ISSN 01691015. doi: 10.1163/156856897X00357. URL http://www.psych.ucsb.edu/.

Diana Bruno and Sofia Schurmann Vignaga. Addenbrooke’s cognitive examination III in the diagnosis of dementia: a critical review. Neuropsychiatric Disease and Treatment, 15: 441, 2019. doi: 10.2147/NDT.S151253. URL https:///pmc/articles/PMC6387595//pmc/articles/PMC6387595/?report=abstract https://www.ncbi.nlm.nih.gov/pmc/articles/PMC6387595/.

Chunting Cai, Chenxi Huang, Chenhui Yang, Haijie Lu, Xin Hong, Fujia Ren, Dan Hong, and Eyk Ng. Altered Patterns of Functional Connectivity and Causal Connectivity in Salience Subnetwork of Subjective Cognitive Decline and Amnestic Mild Cognitive Impairment. Frontiers in Neuroscience, 0:288, 4 2020. ISSN 1662-453X. doi: 10.3389/FNINS.2020.00288.

Cameron J Camp. I am curious-grey: Information seeking and depression across the adult lifespan. Educational Gerontology, 12(4):375–384, 1986. ISSN 15210472. doi: 10.1080/0380127860120412. URL https://www.tandfonline.com/action/journalInformation?journalCode=uedg20.

Nicholas R. Carleton, Myriah K. Mulvogue, Michel A. Thibodeau, Randi E. McCabe, Martin M. Antony, and Gordon J.G. Asmundson. Increasingly certain about uncertainty: Intolerance of uncertainty across anxiety and depression. Journal of Anxiety Disorders, 26(3):468–479, 4 2012. ISSN 08876185. doi: 10.1016/j.janxdis.2012.01.011.

R. Nicholas Carleton. Fear of the unknown: One fear to rule them all?, 2 2016. ISSN 18737897.

L. J. Chang, T. Yarkoni, M. W. Khaw, and A. G. Sanfey. Decoding the Role of the Insula in Human Cognition: Functional Parcellation and Large-Scale Reverse Inference. Cerebral Cortex, 23(3):739–749, 3 2013. ISSN 1047-3211. doi: 10.1093/cercor/bhs065. URL https://academic.oup.com/cercor/article-lookup/doi/10.1093/cercor/bhs065.

Luke Clark, Trevor W. Robbins, Karen D. Ersche, and Barbara J. Sahakian. Reflection Impulsivity in Current and Former Substance Users. Biological Psychiatry, 60(5):515–522, 9 2006. ISSN 00063223. doi: 10.1016/j.biopsych.2005.11.007.

A. D. Craig. How do you feel - now? The anterior insula and human awareness, 1 2009. ISSN 1471003X. URL www.nature.com/reviews/neuro.

A. D. Craig, K. Chen, D. Bandy, and E. M. Reiman. Thermosensory activation of insular cortex. Nature Neuroscience, 3(2):184–190, 2 2000. ISSN 10976256. doi: 10.1038/72131. URL http://neurosci.nature.com.

Hugo D. Critchley, Stefan Wiens, Pia Rotshtein, Arne Öhman, and Raymond J. Dolan. Neural systems supporting interoceptive awareness. Nature Neuroscience 2004 7:2, 7(2):189–195, 1 2004. ISSN 1546-1726. doi: 10.1038/nn1176. URL https://www.nature.com/articles/nn1176.

Ian J. Deary, Janie Corley, Alan J. Gow, Sarah E. Harris, Lorna M. Houlihan, Riccardo E. Marioni, Lars Penke, Snorri B. Rafnsson, and John M. Starr. Age-associated cognitive decline. British Medical Bulletin, 92(1):135–152, 12 2009. ISSN 0007-1420. doi: 10.1093/BMB/LDP033. URL https://academic.oup.com/bmb/article/92/1/135/332828.

Thomas J. DiCiccio and Bradley Efron. Bootstrap confidence intervals. https://doi.org/10.1214/ss/1032280214, 11(3):189–228, 9 1996. ISSN 0883-4237. doi: 10.1214/SS/1032280214. URL https://projecteuclid.org/journals/statistical-science/volume-11/issue-3/Bootstrap-confidence-intervals/10.1214/ss/1032280214.full https://projecteuclid.org/journals/statistical-science/volume-11/issue-3/Bootstrap-confidence-intervals/10.1214/ss/1032280214.short.

Marwa Elamin, Guy Holloway, Thomas H. Bak, and Suvankar Pal. The Utility of the Addenbrooke’s Cognitive Examination Version Three in Early-Onset Dementia. Dementia and Geriatric Cognitive Disorders, 41(1-2):9–15, 3 2016. ISSN 1420-8008. doi: 10.1159/000439248. URL https://www.karger.com/Article/FullText/439248https://www.karger.com/Article/Abstract/439248.

Manes F, Paradiso S, and Robinson RG. Neuropsychiatric effects of insular stroke. The Journal of nervous and mental disease, 187(12):707–712, 1999. ISSN 0022-3018. doi: 10.1097/00005053-199912000-00001. URL https://pubmed.ncbi.nlm.nih.gov/10665464/.

Lisa M. Fitzgerald, Mahnaz Arvaneh, and Paul M. Dockree. Domain-specific and domain-general processes underlying metacognitive judgments. Consciousness and Cognition, 49:264–277, 3 2017. ISSN 1053-8100. doi: 10.1016/J.CONCOG.2017.01.011.

Aikaterini Fotopoulou, Simone Pernigo, Rino Maeda, Anthony Rudd, and Michael A. Kopelman. Implicit awareness in anosognosia for hemiplegia: unconscious interference without conscious re-representation. Brain, 133(12):3564–3577, 12 2010. ISSN 0006-8950. doi: 10.1093/BRAIN/AWQ233. URL https://academic.oup.com/brain/article/133/12/3564/304941.

Christopher Gagne, Ondrej Zika, Peter Dayan, and Sonia J. Bishop. Impaired adaptation of learning to contingency volatility in internalizing psychopathology. eLife, 9:1–51, 12 2020. ISSN 2050084X. doi: 10.7554/ELIFE.61387.

Jacqueline Gottlieb and Pierre-Yves Oudeyer. Towards a neuroscience of active sampling and curiosity. Nature Reviews Neuroscience 2018 19:12, 19(12):758–770, 11 2018. ISSN 1471-0048. doi: 10.1038/s41583-018-0078-0. URL https://www.nature.com/articles/s41583-018-0078-0.

Jeffery Gray and Neil McNaughton. Neuropsychology of anxiety: An enquiry into the functions of septohippocampal theories. Oxford University Press, (1982):469–534, 2003. ISSN 0140-525X. doi: 10.1017/S0140525X00013170.

Dan W. Grupe and Jack B. Nitschke. Uncertainty and anticipation in anxiety: An integrated neurobiological and psychological perspective, 7 2013. ISSN 1471003X. URL www.nature.com/reviews/neuro.

Yuanyuan Gu, Simeng Gu, Yi Lei, and Hong Li. From uncertainty to anxiety: How uncertainty fuels anxiety in a process mediated by intolerance of uncertainty, 2020. ISSN 16875443.

Caroline N. Harada, Marissa C. Natelson Love, and Kristen L. Triebel. Normal Cognitive Aging. Clinics in geriatric medicine, 29(4):737, 11 2013. ISSN 07490690. doi: 10.1016/J.CGER.2013.07.002. URL /pmc/articles/PMC4015335/https://www.ncbi.nlm.nih.gov/pmc/articles/PMC4015335/.

L. M. Harrison, A. Duggins, and K. J. Friston. Encoding uncertainty in the hippocampus. Neural Networks, 19(5):535–546, 6 2006. ISSN 08936080. doi: 10.1016/j.neunet.2005.11.002. URL https://pubmed.ncbi.nlm.nih.gov/16527453/.

Catherine A. Hartley and Elizabeth A. Phelps. Anxiety and decision-making, 7 2012. ISSN 00063223.

Tobias U. Hauser, Michael Moutoussis, Reto Iannaccone, Silvia Brem, Susanne Walitza, Renate Drechsler, Peter Dayan, and Raymond J. Dolan. Increased decision thresholds enhance information gathering performance in juvenile Obsessive-Compulsive Disorder (OCD). PLOS Computational Biology, 13(4):e1005440, 4 2017. ISSN 1553-7358. doi: 10.1371/JOURNAL.PCBI.1005440. URL https://journals.plos.org/ploscompbiol/article?id=10.1371/journal.pcbi.1005440.

Lorraine Hildebrand-Saints and Gifford Weary. Depression and Social Information Gathering. Personality and Social Psychology Bulletin, 15(2):150–160, 6 1989. ISSN 0146-1672. doi: 10.1177/0146167289152002. URL https://journals.sagepub.com/doi/10.1177/0146167289152002http://journals.sagepub.com/doi/10.1177/0146167289152002.

Nikki L. Hill, Jacqueline Mogle, Rachel Wion, Elizabeth Munoz, Nicole DePasquale, Andrea M. Yevchak, and Jeanine M. Parisi. Subjective cognitive impairment and affective symptoms: A systematic review, 12 2016. ISSN 17585341. URL /pmc/articles/PMC5181393//pmc/articles/PMC5181393/?report=abstracthttps://www.ncbi.nlm.nih.gov/pmc/articles/PMC5181393/.

Timothy J. Hohman, Lori L. Beason-Held, and Susan M. Resnick. Cognitive complaints, depressive symptoms, and cognitive impairment: Are they related? Journal of the American Geriatrics Society, 59(10):1908–1912, 10 2011. ISSN 00028614. doi: 10.1111/j.1532-5415.2011.03589.x.

Robert Howard. Subjective cognitive decline: what is it good for? The Lancet Neurology, 19(3):203–204, 3 2020. ISSN 1474-4422. doi: 10.1016/S1474-4422(20)30002-8. URL http://www.thelancet.com/article/S1474442220300028/fulltext http://www.thelancet.com/article/S1474442220300028/abstract https://www.thelancet.com/journals/laneur/article/PIIS1474-4422(20)30002-8/abstract.

Sharpley Hsieh, Samantha Schubert, Christopher Hoon, Eneida Mioshi, and John R. Hodges. Validation of the Addenbrooke’s Cognitive Examination III in Frontotemporal Dementia and Alzheimer’s Disease. Dementia and Geriatric Cognitive Disorders, 36(3-4):242–250, 2013. ISSN 1420-8008. doi: 10.1159/000351671. URL https://www.karger.com/Article/FullText/351671https://www.karger.com/Article/Abstract/351671.

Ryan J. Jacoby, Jonathan S. Abramowitz, Benjamin E. Buck, and Laura E. Fabricant. How is the Beads Task related to intolerance of uncertainty in anxiety disorders? Journal of Anxiety Disorders, 28(6): 495–503, 8 2014. ISSN 18737897. doi: 10.1016/j.janxdis.2014.05.005.

Frank Jessen, Rebecca E. Amariglio, Martin Van Boxtel, Monique Breteler, Mathieu Ceccaldi, Gaël Chételat, Bruno Dubois, Carole Dufouil, Kathryn A. Ellis, Wiesje M. Van Der Flier, Lidia Glodzik, Argonde C. Van Harten, Mony J. De Leon, Pauline McHugh, Michelle M. Mielke, Jose Luis Molinuevo, Lisa Mosconi, Ricardo S. Osorio, Audrey Perrotin, Ronald C. Petersen, Laura A. Rabin, Lorena Rami, Barry Reisberg, Dorene M. Rentz, Perminder S. Sachdev, Vincent De La Sayette, Andrew J. Saykin, Philip Scheltens, Melanie B. Shulman, Melissa J. Slavin, Reisa A. Sperling, Robert Stewart, Olga Uspenskaya, Bruno Vellas, Pieter Jelle Visser, and Michael Wagner. A conceptual framework for research on subjective cognitive decline in preclinical Alzheimer’s disease. Alzheimer’s and Dementia, 10(6):844–852, 11 2014. ISSN 15525279. doi: 10.1016/j.jalz.2014.01.001 URL http://dx.doi.org/10.1016/j.jalz.2014.01.001.

Frank Jessen, Rebecca E. Amariglio, Rachel F. Buckley, Wiesje M. van der Flier, Ying Han, José Luis Molinuevo, Laura Rabin, Dorene M. Rentz, Octavio Rodriguez-Gomez, Andrew J. Saykin, Sietske A.M. Sikkes, Colette M. Smart, Steffen Wolfsgruber, and Michael Wagner. The character-isation of subjective cognitive decline, 3 2020. ISSN 14744465. URL https://doi.org/10.1016/.

Thomas E Joiner, Gerald I Metalsky, Jennifer Katz, and Steven R.H. Beach. Depression and excessive reassurance-seeking. Psychological Inquiry, 10(4):269–278, 1999. ISSN 1047840X. doi: 10.1207/s15327965pli1004{\}1. URL https://www.tandfonline.com/action/journalInformation?journalCode=hpli20.

Peter R Jones, Linnea Landin, Aisha McLean, Mordechai Z. Juni, Laurence T Maloney, Marko Nardini, and Tessa M Dekker. Efficient visual information sampling develops late in childhood. Journal of Experimental Psychology: General, 148(7):1138–1152, 7 2019. ISSN 1939-2222. doi: 10.1037/xge0000629. URL http://supp.apa.org/psycarticles/supplemental/xge0000629/xge0000629_supp.htmlhttp://doi.apa.org/getdoi.cfm?doi=10.1037/xge0000629.

Mordechai Z. Juni, Todd M. Gureckis, and Laurence T. Maloney. Information sampling behavior with explicit sampling costs. Decision, 3(3):147–168, 2016. ISSN 23259973. doi: 10.1037/dec0000045.

Hans-Otto Karnath, Bernhard Baier, and Thomas Nägele. Awareness of the Functioning of One’s Own Limbs Mediated by the Insular Cortex? Journal of Neuroscience, 25(31):7134–7138, 8 2005. ISSN 0270-6474. doi: 10.1523/JNEUROSCI.1590-05.2005. URL https://www.jneurosci.org/content/25/31/7134https://www.jneurosci.org/content/25/31/7134.abstract.

M Kleiner, D Brainard, Denis Pelli, A Ingling, R Murray, and C Broussard. What’s new in psychtoolbox-3. Perception, 36(14):1–16, 2007. ISSN 0301-0066. URL https://nyuscholars.nyu.edu/en/publications/whats-new-in-psychtoolbox-3.

Richard J. Kryscio, Erin L. Abner, Gregory E. Cooper, David W. Fardo, Gregory A. Jicha, Peter T. Nelson, Charles D. Smith, Linda J. Van Eldik, Lijie Wan, and Frederick A. Schmitt. Self-reported memory complaints. Neurology, 83(15):1359–1365, 10 2014. ISSN 0028-3878. doi: 10.1212/WNL.0000000000000856. URL https://n.neurology.org/content/83/15/1359 https://n.neurology.org/content/83/15/1359.abstract.

William B. Locander and Peter W. Hermann. The Effect of Self-Confidence and Anxiety on Information Seeking in Consumer Risk Reduction. Journal of Marketing Research, 16(2):268, 5 1979. ISSN 00222437. doi: 10.2307/3150690. URL https://www.jstor.org/stable/3150690?origin=crossref.

Victoria C. Martin, Daniel L. Schacter, Michael C. Corballis, and Donna Rose Addis. A role for the hippocampus in encoding simulations of future events. Proceedings of the National Academy of Sciences of the United States of America, 108(33):13858–13863, 8 2011. ISSN 00278424. doi: 10.1073/PNAS.1105816108/-/DCSUPPLEMENTAL. URL https://www.pnas.org/content/108/33/13858https://www.pnas.org/content/108/33/13858.abstract.

Peter M. McEvoy and Alison E.J. Mahoney. Achieving certainty about the structure of intolerance of uncertainty in a treatment-seeking sample with anxiety and depression. Journal of Anxiety Disorders, 25(1):112–122, 1 2011. ISSN 0887-6185. doi: 10.1016/J.JANXDIS.2010.08.010.

Laura McWhirter, Craig Ritchie, Jon Stone, and Alan Carson. Functional cognitive disorders: a systematic review. The Lancet Psychiatry, 7(2):191–207, 2 2020. ISSN 2215-0366. doi: 10.1016/S2215-0366(19)30405-5.

Marcelo D. Mendonça, Luísa Alves, and Paulo Bugalho. From Subjective Cognitive Complaints to Dementia, 3 2016. ISSN 19382731. URL https://journals.sagepub.com/doi/10.1177/1533317515592331?url_ver=Z39.88-2003&rfr_id=ori%3Arid%3Acrossref.org&rfr_dat=cr_pub++0pubmed.

Mark J. Millan, Yves Agid, Martin Brüne, Edward T. Bullmore, Cameron S. Carter, Nicola S. Clayton, Richard Connor, Sabrina Davis, Bill Deakin, Robert J. Derubeis, Bruno Dubois, Mark A. Geyer, Guy M. Goodwin, Philip Gorwood, Thérèse M. Jay, Marian Joëls, Isabelle M. Mansuy, Andreas Meyer-Lindenberg, Declan Murphy, Edmund Rolls, Bernd Saletu, Michael Spedding, John Sweeney, Miles Whittington, and Larry J. Young. Cognitive dysfunction in psychiatric disorders: Characteristics, causes and the quest for improved therapy, 2 2012. ISSN 14741776. URL www.nature.com/reviews/drugdisc.

Jayne Morriss, Martin Gell, and Carien M. van Reekum. The uncertain brain: A co-ordinate based meta-analysis of the neural signatures supporting uncertainty during different contexts, 1 2019. ISSN 18737528.

Ho Namkung, Sun Hong Kim, and Akira Sawa. The Insula: An Underestimated Brain Area in Clinical Neuroscience, Psychiatry, and Neurology, 4 2017. ISSN 1878108X.

Thomas O. Nelson. Metamemory: A Theoretical Framework and New Findings. Psychology of Learning and Motivation - Advances in Research and Theory, 26(C):125–173, 1 1990. ISSN 0079-7421. doi: 10.1016/S0079-7421(08)60053-5.

Martin P. Paulus and Murray B. Stein. An Insular View of Anxiety, 8 2006. ISSN 00063223.

Ivanna M. Pavisic, Kirsty Lu, Sarah E. Keuss, Sarah-Naomi James, Christopher A. Lane, Thomas D. Parker, Ashvini Keshavan, Sarah M. Buchanan, Heidi Murray-Smith, David M. Cash, William Coath, Andrew Wong, Nick C. Fox, Sebastian J. Crutch, Marcus Richards, and Jonathan M. Schott. Subjective cognitive complaints at age 70: associations with amyloid and mental health. Journal of Neurology, Neurosurgery & Psychiatry, 0:2020–325620, 5 2021. ISSN 0022-3050. doi: 10.1136/jnnp-2020-325620. URL https://jnnp.bmj.com/lookup/doi/10.1136/jnnp-2020-325620.

Pierre Petitet, Bahaaeddin Attaallah, Sanjay G Manohar, and Masud Husain. The computational cost of active information sampling before decision-making under uncertainty. Nature Human Behaviour, pages 1–12, 5 2021. doi: 10.1038/s41562-021-01116-6. URL https://doi.org/10.1038/s41562-021-01116-6.

L. D. Phillips, W. L. Hays, and W. Edwards. Conservatism in Complex Probabilistic Inference. IEEE Transactions on Human Factors in Electronics, HFE-7(1):7–18, 1966. ISSN 21682852. doi: 10.1109/THFE.1966.231978.

Diego A. Pizzagalli, Allison L. Jahn, and James P. O’Shea. Toward an objective characterization of an anhedonic phenotype: A signal-detection approach. Biological Psychiatry, 57(4):319–327, 2 2005. ISSN 0006-3223. doi: 10.1016/J.BIOPSYCH.2004.11.026.

Diego A. Pizzagalli, Dan Iosifescu, Lindsay A. Hallett, Kyle G. Ratner, and Maurizio Fava. Reduced hedonic capacity in major depressive disorder: Evidence from a probabilistic reward task. Journal of Psychiatric Research, 43(1):76–87, 11 2008. ISSN 0022-3956. doi: 10.1016/J.JPSYCHIRES.2008.03.001.

Erdem Pulcu and Michael Browning. The Misestimation of Uncertainty in Affective Disorders. Trends in Cognitive Sciences, 8 2019. ISSN 1364-6613. doi: 10.1016/J.TICS.2019.07.007. URL https://www.sciencedirect.com/science/article/pii/S1364661319301822.

Louise M. Reid and Alasdair M.J. MacLullich. Subjective memory complaints and cognitive impairment in older people, 11 2006. ISSN 14208008. URL www.karger.comwww.karger.com/.

Francesco Rigoli, Jochen Michely, Karl J. Friston, and Raymond J. Dolan. The role of the hippocampus in weighting expectations during inference under uncertainty. Cortex, 115:1–14, 6 2019. ISSN 0010-9452. doi: 10.1016/J.CORTEX.2019.01.005. URL https://www.sciencedirect.com/science/article/pii/S0010945219300206#fig1.

Kevin G. Saulnier, Nicholas P. Allan, Amanda M. Raines, and Norman B. Schmidt. Depression and Intolerance of Uncertainty: Relations between Uncertainty Subfactors and Depression Dimensions. https://doi.org/10.1080/00332747.2018.1560583, 82(1):72–79, 1 2019. doi: 10.1080/00332747.2018.1560583. URL https://www.tandfonline.com/doi/abs/10.1080/00332747.2018.1560583.

Daniel L. Schacter, Donna Rose Addis, and Randy L. Buckner. Episodic Simulation of Future Events. Annals of the New York Academy of Sciences, 1124(1):39–60, 3 2008. ISSN 1749-6632. doi: 10.1196/ANNALS.1440.001. URL https://onlinelibrary.wiley.com/doi/full/10.1196/annals.1440.001 https://onlinelibrary.wiley.com/doi/abs/10.1196/annals.1440.001 https://nyaspubs.onlinelibrary.wiley.com/doi/10.1196/annals.1440.001.

Daniel L Schacter, Donna Rose Addis, and Karl K Szpunar. Escaping the Past: Contributions of the Hippocampus to Future Thinking and Imagination. 2017. doi: 10.1007/978-3-319-50406-3{\}14.

Tali Sharot and Cass R. Sunstein. How people decide what they want to know. Nature Human Behaviour 2020 4:1, 4(1):14–19, 1 2020. ISSN 2397-3374. doi: 10.1038/s41562-019-0793-1. URL https://www.nature.com/articles/s41562-019-0793-1.

Tania Singer, Hugo D. Critchley, and Kerstin Preuschoff. A common role of insula in feelings, empathy and uncertainty. Trends in cognitive sciences, 13(8):334–340, 8 2009. ISSN 1364-6613. doi: 10.1016/J.TICS.2009.05.001. URL https://pubmed.ncbi.nlm.nih.gov/19643659/.

Lucia Spinazzola, Lorenzo Pia, Alessia Folegatti, Clelia Marchetti, and Anna Berti. Modular structure of awareness for sensorimotor disorders: Evidence from anosognosia for hemiplegia and anosognosia for hemianaesthesia. Neuropsychologia, 46(3):915–926, 1 2008. ISSN 0028-3932. doi: 10.1016/J.NEUROPSYCHOLOGIA.2007.12.015.

Bryan A. Strange, Andrew Duggins, William Penny, Raymond J. Dolan, and Karl J. Friston. Information theory, novelty and hippocampal responses: Unpredicted or unpredictable? Neural Networks, 18(3):225–230, 4 2005. ISSN 08936080. doi: 10.1016/j.neunet.2004.12.004.

Ema Tanovic, Dylan G. Gee, and Jutta Joormann. Intolerance of uncertainty: Neural and psychophysiological correlates of the perception of uncertainty as threatening, 3 2018. ISSN 18737811.

Dustin Tingley, Teppei Yamamoto, Kentaro Hirose, Luke Keele, and Kosuke Imai. Mediation: R package for causal mediation analysis. Journal of Statistical Software, 59(5): 1–38, 8 2014. ISSN 15487660. doi: 10.18637/jss.v059.i05. URL https://www.jstatsoft.org/index.php/jss/article/view/v059i05/v59i05.pdf https://www.jstatsoft.org/index.php/jss/article/view/v059i05.

Michael J. Tobia, Vittorio Iacovella, Ben Davis, and Uri Hasson. Neural systems mediating recognition of changes in statistical regularities. NeuroImage, 63(3):1730–1742, 11 2012. ISSN 10538119. doi: 10.1016/j.neuroimage.2012.08.017. URL https://pubmed.ncbi.nlm.nih.gov/22906790/.

Lucina Q. Uddin, Jason S. Nomi, Benjamin Hébert-Seropian, Jimmy Ghaziri, and Olivier Boucher. Structure and Function of the Human Insula, 7 2017. ISSN 15371603. URL /pmc/articles/PMC6032992//pmc/articles/PMC6032992/?report=abstracthttps://www.ncbi.nlm.nih.gov/pmc/articles/PMC6032992/.

Sturm VE, Rosen HJ, Allison S, Miller BL, and Levenson RW. Self-conscious emotion deficits in frontotemporal lobar degeneration. Brain : a journal of neurology, 129(Pt 9):2508–2516, 9 2006. ISSN 1460-2156. doi: 10.1093/BRAIN/AWL145. URL https://pubmed.ncbi.nlm.nih.gov/16844714/https://pubmed.ncbi.nlm.nih.gov/16844714/?dopt=Abstract.

Julia A. Weiler, Boris Suchan, and Irene Daum. Foreseeing the future: occurrence probability of imagined future events modulates hippocampal activation. Hippocampus, 20(6):685–690, 6 2010. ISSN 1098-1063. doi: 10.1002/HIPO.20695. URL https://pubmed.ncbi.nlm.nih.gov/19693779/https://pubmed.ncbi.nlm.nih.gov/19693779/?dopt=Abstract.

Susan Whitfield-Gabrieli and Alfonso Nieto-Castanon. Conn: A Functional Connectivity Toolbox for Correlated and Anticorrelated Brain Networks. https://home.liebertpub.com/brain, 2(3):125–141, 8 2012. doi: 10.1089/BRAIN.2012.0073. URL https://www.liebertpub.com/doi/abs/10.1089/brain.2012.0073.

A. S. Zigmond and R. P. Snaith. The Hospital Anxiety and Depression Scale. Acta Psychiatrica Scandinavica, 67(6):361–370, 1983. doi: 10.1111/J.1600-0447.1983.TB09716.X.

